# Using transition density models to interpret experimental optical spectra of exciton-coupled cyanine (iCy3)_2_ dimer probes of local DNA conformations at or near functional protein binding sites

**DOI:** 10.1101/2023.08.26.554948

**Authors:** Dylan Heussman, Lulu Enkhbaatar, Mohammed I. Sorour, Kurt A. Kistler, Peter H. von Hippel, Spiridoula Matsika, Andrew H. Marcus

## Abstract

Exciton-coupled chromophore dimers are an emerging class of optical probes for studies of site-specific biomolecular interactions. Applying accurate theoretical models for the electrostatic coupling of a molecular dimer probe is a key step for simulating its optical properties and analyzing spectroscopic data. In this work, we compare experimental absorbance and circular dichroism (CD) spectra of ‘internally-labeled’ (iCy3)_2_ dimer probes inserted site-specifically into DNA fork constructs to theoretical calculations of the structure and geometry of these exciton-coupled dimers. We compare transition density models of varying levels of approximation to determine conformational parameters of the (iCy3)_2_ dimer-labeled DNA fork constructs. By applying an atomistically detailed transition charge (TQ) model, we can distinguish between dimer conformations in which the stacking and tilt angles between planar iCy3 monomers are varied. A major strength of this approach is that the local conformations of the (iCy3)_2_ dimer probes that we determined can be used to infer information about the structures of the DNA framework immediately surrounding the probes at various positions within the constructs, both deep in the duplex DNA sequences and at sites at or near the DNA fork junctions where protein complexes bind to discharge their biological functions.

## 1. Introduction

The biological properties of nucleic acids rely on their ability to undergo both base-sequence-specific and non-base-sequence-specific interactions with proteins, which play a major role in guiding their assembly into multi-subunit protein-nucleic acid complexes. Understanding the physical-chemical basis of these interactions is an area of active research. For example, understanding in structural and dynamical terms how local conformational fluctuations of the nucleobases and sugar-phosphate backbones of DNA facilitate binding to proteins can provide key insights about the functional mechanisms of macromolecular machines, as discussed at length in a recent paper from our laboratory [1]. Although established structural tools such as NMR and X-ray crystallography provide detailed molecular-level information about stable protein-nucleic acid complexes at high concentrations [2, 3], such methods are not well-suited to probe site-specific protein-DNA interactions at the micromolar-to-nanomolar concentrations at which these complexes typically assemble and function.

An approach that is well suited for studies at physiological concentrations is to perform spectroscopic experiments on a DNA molecule that has been site-specifically labeled with fluorescent optical probes [4-6]. For example, the monomer forms of the fluorescent cyanine dyes Cy3 and Cy5 can be chemically attached to a nucleic acid base or to the distal end of a nucleic acid chain using a flexible linker. Such fluorescently labeled DNA constructs are commercially available for fluorescence microscopy, gene sequencing technologies, and other bio-analytical applications. Measurements based on Förster resonant energy transfer (FRET) are regularly used to determine structural changes of a DNA strand labeled with a Cy3 ‘donor’ and a Cy5 ‘acceptor’ on the length scale of a few nanometers [7-11]. Alternatively, pairs of Cy3 dye molecules can be attached ‘internally’ (termed ‘iCy3’) and rigidly at defined opposing positions within the sugar-phosphate backbones of complementary DNA strands to create an exciton-coupled (iCy3)_2_ dimer-probe-labeled single-stranded (ss) – double-stranded (ds) DNA construct, in which the dimer probes are site-specifically inserted relative to the fork junction (see Fig. 1) [12-15]. DNA constructs, including model Holliday junctions used in recombination, have also been studied using exciton-coupled dimers of Cy5 [16-19]. Such approaches have the additional virtue of providing *local* structural information at defined and functionally important positions within the DNA frameworks of active protein-DNA complexes.

**Figure 1.**
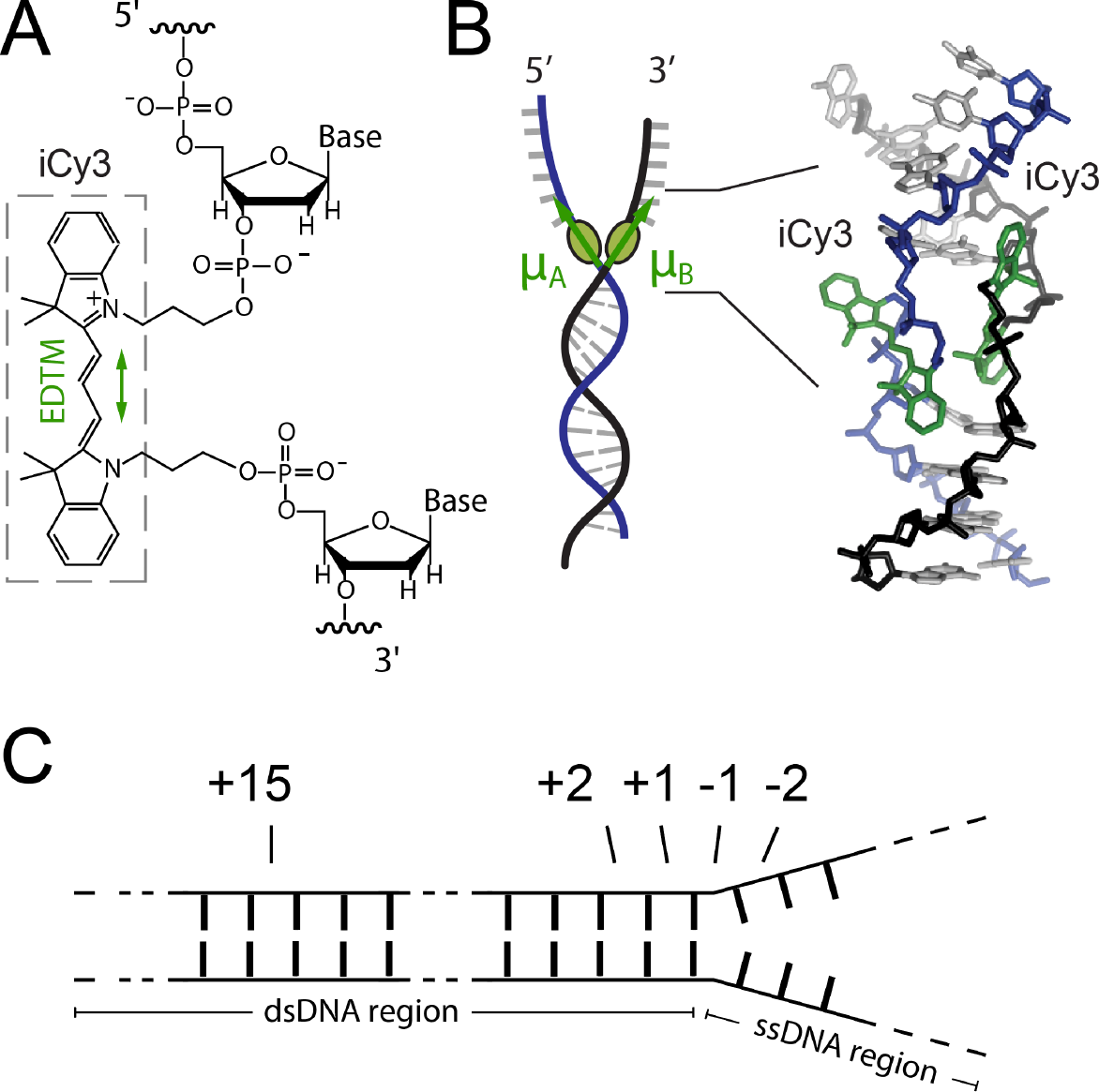
(*A*) The Lewis structure of the iCy3 chromophore is shown with its 3’ and 5’ linkages to the sugar-phosphate backbone of a local segment of ssDNA. The electric dipole transition moment (EDTM, indicated by green double-headed arrow) is orientated parallel to the all-trans trimethine bridge of the iCy3 chromophore. (*B*) An (iCy3)2 dimer-labeled DNA fork construct contains the dimer probe near the ss-dsDNA fork junction. The local secondary structure of the sugar-phosphate backbones at the probe insertion site is reflected by the dimer probe conformation. The sugar-phosphate backbones of the conjugate DNA strands are shown in black and blue, the bases are shown in gray, and the iCy3 chromophores are shown in green. (*C*) The position of the dimer probe is indicated relative to the pseudo-fork junction using positive (negative) integers in the direction toward the double-(single-) stranded region.

An internally-attached iCy3 within a DNA single-strand acts as a molecular bridge between flanking bases and as an extension of the sugar-phosphate backbone between adjacent nucleotides (Fig. 1*A*) [6]. The linear absorbance spectrum of free Cy3 monomer in polar solvent, as well as when it is attached internally as an iCy3 monomer in a nucleic acid framework, is relatively insensitive to the local environment of the chromophore and exhibits a pronounced vibronic progression [13, 20, 21]. By annealing two complementary strands of DNA with opposed iCy3 labeling positions, an exciton-coupled (iCy3)_2_ dimer probe (with monomers labeled *A* and *B*) can be formed at a predetermined position within a model DNA fork construct (see Fig. 1*B*) and its optical spectrum is sensitive to the dimer conformation, which can thus serve to provide information about the geometry and stability of the adjacent DNA segments. The annealed DNA constructs that we use in this study contain both dsDNA and ssDNA regions, and the (iCy3)_2_ dimer probe can be selectively inserted at varying positions relative to the ss-dsDNA fork junction (see Fig. 1*C* for probe-labeling nomenclature). The closely-spaced monomers of the (iCy3)_2_ dimer probe can couple through an electrostatic (exciton-coupled) interaction *J* [13-15]. The value of *J* and the resulting spectroscopic properties of the (iCy3)_2_ dimer-labeled DNA construct depend sensitively on the relative orientation and spacing between the iCy3 monomers on the length scale of a few Angstroms. Such cyanine dimer-labeled DNA constructs have been used, for example, to study electronically excited-state dynamics in vibronically coupled dimers and higher-order chromophore networks [16-18, 22, 23].

The value of *J* can be calculated using quantum chemical models of the monomer electronic transition density [24-26]. In previous work, we studied the mean local conformations and conformational disorder of (iCy3)_2_ dimer-probe-labeled ss-dsDNA fork constructs for specific probe insertion-site positions. In these studies, we compared experimental absorbance, circular dichroism (CD) and two-dimensional fluorescence spectroscopy (2DFS) to theoretically-derived spectra in which we applied the ‘point-dipole’ (PD) [13] and the ‘extended-dipole’ (ED) [14, 15] models to efficiently approximate *J*. For intermolecular separations smaller than the iCy3 monomer dimension, the PD model unrealistically assumes that the transition density can be approximated as a point-dipole located at the molecular center-of-mass (see Fig. 2*A*). The finite dimension of the molecule can be further accounted for by using the ED model, which represents the transition density as a line segment that is oriented parallel to the transition dipole moment with equal and opposite charges on its endpoints (Fig. 2*B*). While the results of our prior studies suggested that the PD and ED models can provide self-consistent and reliable structural interpretations of optical spectra for (iCy3)_2_ dimer-labeled ss-dsDNA constructs, a firm understanding of the strengths and limitations of these models has not been fully established. In the current work, we apply an atomistically-detailed electrostatic coupling model that is based on ‘transition charges’ (TQs) derived from the *ab initio* transition density to compare to the optical spectra of (iCy3)_2_ dimer-labeled ss-dsDNA constructs. This TQ model was developed previously to describe more realistically, in comparison to the PD and ED models, the transition densities of interacting iCy3 monomers by assigning individual transition electrostatic potential (trESP) charges to the atomic coordinates [27] (Fig. 2*C*). The trESP method uses an approach like that for deriving ground electronic state charges, except that the electrostatic potential is obtained from the transition density matrix between ground and excited electronic states while neglecting the nuclear contributions to the electrostatic potential. The resulting trESP surface is subsequently coarse-grained and decomposed into transition point charges centered on the atomic coordinates of the Cy3 molecule.

**Figure 2.**
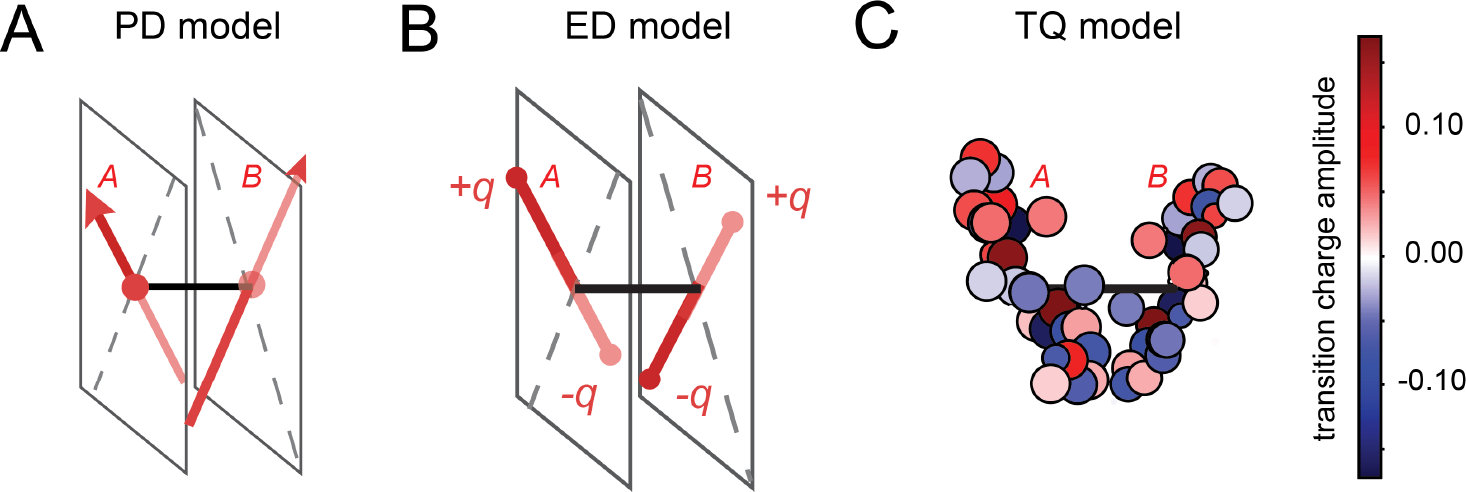
Illustration of transition charge density models for the estimation of the electrostatic coupling of (iCy3)_2_ dimer-labeled ss-dsDNA constructs. (*A*) Point-dipole (PD) model. (*B*) Extended-dipole (ED) model. (*C*) Transition charge (TQ) density model. See text for further explanation.

To characterize the conformations of (iCy3)_2_ dimer-labeled ss-dsDNA constructs, we have defined a minimum set of structural parameters (see Fig. 3). For the PD and ED models, the conformation is fully specified by the ‘tilt,’ *θ*_*AB*_, ‘twist,’ *ϕ*_*AB*_, and ‘inter-chromophore separation,’

**Figure 3.**
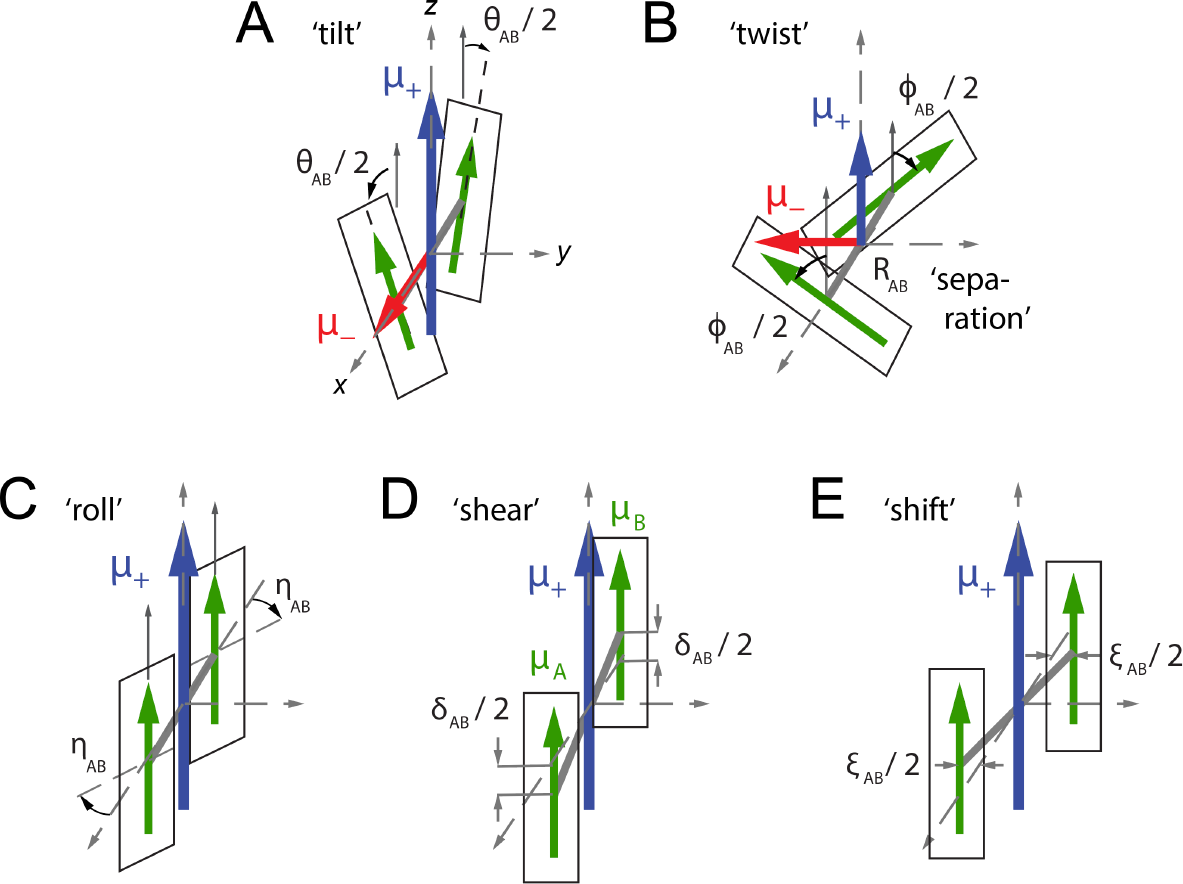
Conformational parameters used for the electrostatic coupling models studied in this work. Green arrows indicate the monomer EDTMs (labeled *A* and *B*), and rectangles indicate the π-conjugation planes of the iCy3 trimethine groups (as shown in Fig. 1*A*). The electrostatic coupling between the iCy3 chromophores gives rise to symmetric (+) and anti-symmetric (–) excitons, which are indicated by blue and red arrows, respectively. (*A*) ‘Tilt’ angle *θ*_*AB*_. (*B*) ‘Twist’ angle *ϕ*_*AB*_ and inter-chromophore center-of-mass separation *R*_*AB*_. (*C*) ‘Roll’ angle *η*_*AB*_ (illustrating a symmetric rotation). (*D*) Vertical ‘shear’ displacement *δ*_*AB*_. (*E*) Horizontal ‘shift’ displacement *ξ*_*AB*_. In panels (*C* – *E*), the tilt and twist angles have been set to zero.

*R*_*AB*_ (Figs. 3*A* – 3*B*). For the TQ model calculations, which add more dimensionality to the model of the transition density, additional structural parameters are needed to distinguish between relative orientations and translational displacements of the iCy3 monomers. We thus defined the ‘roll’ angle, *η*_*AB*_, the vertical ‘shear’ displacement, *δ*_*AB*_, and the horizontal ‘shift’ displacement, *ξ*_*AB*_ (Figs. 3*C* – 3*E*). For a specific set of conformational parameters, and depending on the choice of coupling model, we calculated the value of *J* and simulated the optical spectra, which we compared to previously reported experimental data [15]. We thus obtained ‘optimized values’ for the conformational parameters by applying the above procedure iteratively in combination with a multi-parameter optimization procedure [28, 29].

As we describe in further detail below, we find that the three electrostatic coupling models for the (iCy3)_2_ dimer-labeled ss-dsDNA constructs provide nearly identical results for the twist angle, *ϕ*_*AB*_, and similar results for the tilt, *θ*_*AB*_, roll, *η*_*AB*_ and inter-chromophore separation *R*_*AB*_. In principle, information about the roll angle, *η*_*AB*_, is only available using the atomistically-detailed TQ model. However, by systematically rejecting sterically overlapping dimer conformations (approximating the van der Waals radii of the major component atoms of the iCy3 monomers as *r*_*w*_ ∼1.5Å), corroborating information between the three models for all structural parameters can be obtained. By directly comparing the results of the PD, ED and TQ models, which each represents a different level of approximation for the transition density, we establish the reliability of the approach for determining structural information from optical spectra. Our ultimate intent in developing these methods – in combination with single-molecule spectroscopic measurements – is to interpret and understand the conformations and dynamics of biologically important DNA sites that are positioned immediately adjacent to the (iCy3)_2_ dimer probes and effectively control the structures and dynamics of the dimer probes themselves.

While in the current study we focus primarily on the spectroscopic properties of exciton-coupled (iCy3)_2_ homodimer-labeled DNA constructs, the TQ model could be well applied to calculate the electrostatic coupling between closely spaced donor-acceptor chromophore pairs in Förster resonance energy transfer (FRET) experiments [11, 30]. In such situations, the TQ model would provide a more accurate description of the electrostatic coupling. However, for many experiments in which a donor-acceptor FRET pair is used to site-specifically label a biological macromolecule, the inter-chromophore separation is on the order of the Förster distance of a few nanometers such that the electrostatic coupling is relatively weak. For weakly coupled donor-acceptor FRET pairs, the point-dipole model provides a reasonable approximation of the electrostatic coupling, as indicated by the calculations presented below (see Fig. 6 below).

## II. Materials and Methods

### A. Effective Mode Hamiltonian for (iCy3)_2_ Dimer-Labeled DNA Constructs

The Cy3 monomer consists of a conjugated trimethine bridge, which cojoins two indole-like groups (Fig. 1*A*). The absorbance spectrum of the Cy3 monomer in solution, or when it is attached internally (as iCy3) to DNA, is minimally affected by its environment, and exhibits a pronounced vibronic progression [13]. The trimethine bridge of the Cy3 monomer exists in an *all-trans* configuration, and the lowest energy electronic transition is a π → π^*^ transition [31]. Like many π-conjugated molecules, Cy3 exhibits numerous Franck-Condon-active modes, which range in energy from tens-to-several hundred wavenumbers [32]. Nevertheless, the homogeneous spectral lineshape of the Cy3 monomer in solution is broadened due to rapid electronic dephasing and thermal occupation of low energy vibrational levels. The lowest energy electronic transition from ground state |*g*⟩ to excited state |*e*⟩ is predominantly coupled to one low-frequency (∼30 cm^−1^) symmetric bending vibration along the trimethine bridge, which couples anharmonically to a cluster of relatively high-frequency modes in the vicinity of ∼1,300 cm^-1^ [21]. The spectrum of the Cy3 monomer in solution – and that of the iCy3 monomer in the DNA sugar-phosphate backbone – can be simulated using a relatively simple quantum mechanical Hamiltonian, 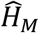, where the |*g*⟩ → |*e*⟩ transition (with energy ε_*eg*_= ∼18,250 cm^-1^) is coupled to a single ‘effective’ harmonic mode (with energy ℏω_0_ = ∼1,100 cm^-1^) [13-15]. The monomer Hamiltonian is written:

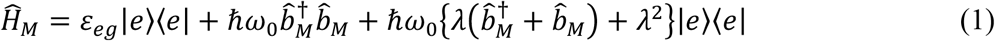

where 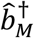 and 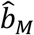 are, respectively, the operators for creating and annihilating a vibrational excitation in the ground and excited electronic potential energy surfaces. These operators obey the commutation relation 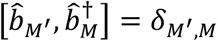, where *δM*^*′*^,*M* is the Kronecker delta function and the monomer labels *M*^′^, *M* ∈ {*A, B*}.

The strength of the vibronic coupling is characterized by Franck-Condon (FC) factors,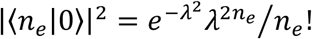, where |0⟩ = |*n*_*g*_ = 0⟩|*g*⟩ is the zero vibrational state of the ground electronic state, and |*n*_*e*_⟩ is the vibrational state of the electronically excited state with vibrational quantum number *n*_*e*_. The FC factors depend on the Huang-Rhys parameter (λ^2^= *d*^2^ω_0_*m*/2ℏ = ∼0.57) where *d* is the displacement of the electronically excited vibrational potential energy surface relative to the ground state surface and *m* is the oscillator reduced mass [33]. The monomer electric dipole transition moment (EDTM) has magnitude μ_*eg*_ = ∼12.8 D and is oriented parallel to the iCy3 trimethine bridge, along the long axis of the molecule (Fig. 1*A*).

For the (iCy3)_2_ dimer probe, the monomer EDTMs can couple to one another through an electrostatic interaction *J*. The Holstein-Frenkel (H-F) Hamiltonian of the coupled dimer is written [24]

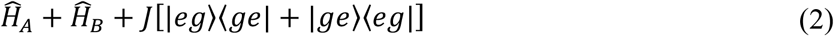

where |*eg*⟩ is the product state in which monomer *A* is electronically excited and monomer *B* is unexcited. For non-zero electrostatic coupling *J*, the absorbance spectrum of the (iCy3)_2_ dimer has a shape roughly like that of the monomer. However, the electrostatic interaction leads to each of the vibronic features (i.e., 0-0, 0-1, …) being split and additionally broadened into symmetric (+) and anti-symmetric (–) sub-bands. The relative peak intensities of the absorbance spectrum depend on – in addition to the Franck-Condon factors that affect the vibronic features of the iCy3 monomer – the local conformation of the dimer, which determines the magnitudes of the symmetric and anti-symmetric transition dipole moments 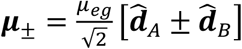 where 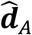 and 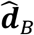 are unit vectors that specify the monomer EDTM directions. The symmetric and anti-symmetric excitons of each consist of a manifold of delocalized electronic-vibrationally coupled states, which are superpositions of electronic-vibrational products of the *A* and *B* monomer states given by 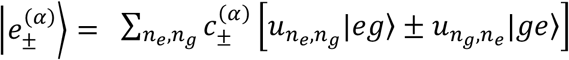[25]. The coefficients 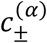 depend on the vibrational coordinates of the monomers, 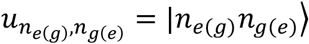 is the vibrational product state, *n*_*e*(*g*)_ is the vibrational quantum number in the electronic excited (ground) state, and α = (0, 1, …) is a state index in order of increasing energy.

The (iCy3)_2_ dimer absorbance spectrum is the sum of symmetric (+) and anti-symmetric (−) excitons

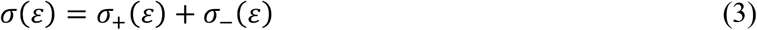

where 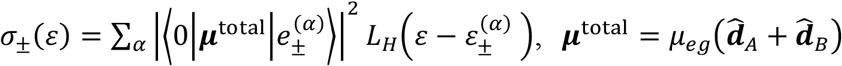 is the collective EDTM and 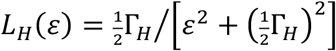 is a Lorentzian homogeneous lineshape representing the transition with eigen-energy 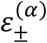 and full-width-at-half-maximum (FWHM) line width Γ_*H*_ (set equal to 186 cm^-1^, as in previous work [13-15]). Similarly, the dimer CD spectrum is the sum of symmetric and anti-symmetric rotational strengths

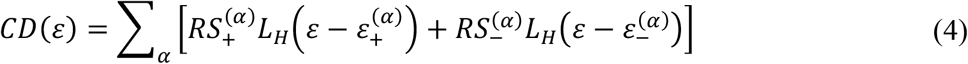

where 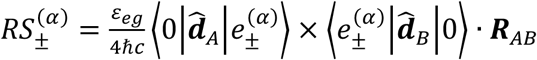. In the above expressions, we have defined the ground vibrational-electronic state of the *AB* dimer |0⟩ = |*n*_*A*_ = 0, *n*_*B*_ = 0⟩|*gg*⟩.

The (iCy3)_2_ dimer conformation may vary from molecule to molecule due to local (and position-specific) DNA ‘breathing’ fluctuations, so that the homogeneous dimer absorbance and CD line shapes are convolved with an inhomogeneous distribution function 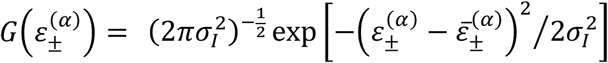, which is centered at the average transition energy 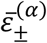 and has standard deviation σ_*A*_. The final expressions for the absorbance and CD spectra are given, respectively, by the Voigt profiles:

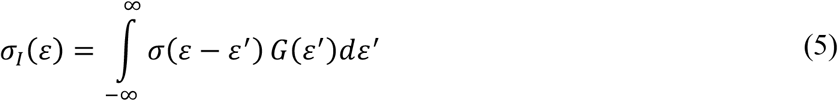

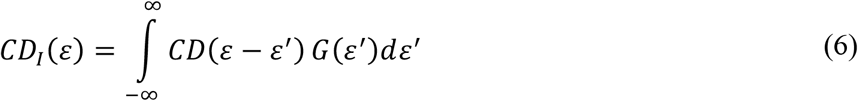

### B. Transition Density Models of the Electrostatic Coupling in (iCy3)_2_ Dimer-Labeled DNA Constructs

For our analyses of experimental absorbance and CD spectra in terms of the (iCy3)_2_ dimer conformation, it was necessary to implement a computationally-efficient means to approximate the electrostatic coupling *J* that appears in Eq. (2). The electrostatic coupling can be modeled in terms of the Coulomb interaction between the monomer transition densities [34],

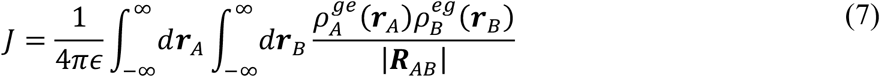

where *R*_*AB*_ = *r*_*B*_ − *r*_*A*_ is the inter-chromophore center-of-mass separation vector, 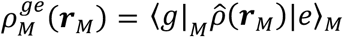 is the monomer transition density matrix element, and ϵ is the electric permittivity. In the point-dipole (PD) approximation, the electrostatic coupling is given by [34],

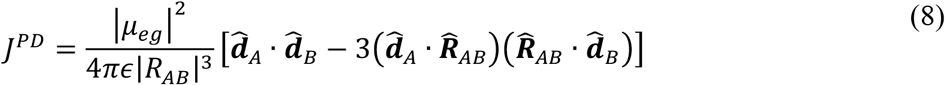

The ‘extended dipole’ (ED) model accounts for the finite size of the iCy3 chromophore by including a one-dimensional displacement vector *l* that lies parallel to the monomer EDTM [35-37].

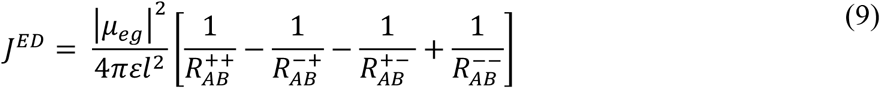

Equation (9) approximates the transition density for each monomer as two separate point charges of equal magnitude (*q*) and opposite sign, which lie separated by the distance *l* such that *ql* = μ_()_ (Fig. 2*B*). The distances between point charges are given by 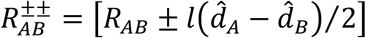 and 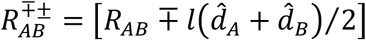. In the transition charge (TQ) model, the electrostatic interaction is approximated as a discrete sum over trESP transition charges (*q*_*i*_), which are assigned to the centers of each atomic coordinate of the monomer.

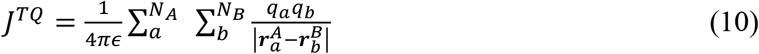

In Eq. (10), 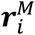 is the position of the *i*^th^ atom of chromophore *M* (∈ {*A, B*}) and *N*_*M*_ is the number of atoms in chromophore *M*. The values of the transition charges *q*_*i*_ were obtained from an *ab initio* quantum chemical calculation of the iCy3 chromophore [27].

To ensure self-consistency between the results of our PD, ED and TQ calculations, we maintained a constant value for the magnitude of the iCy3 EDTM (μ_*eg*_ = ∼12.8 D), which we determined experimentally by integrating the spectral lineshape [13]. Under the PD approximation, the value of μ_*eg*_ is uniquely defined. However, in the ED and TQ models the value of μ_*eg*_ can be scaled by independently varying the transition charges and the inter-charge separations. For example, under the ED model the transition point-charges (*q* = ±0.38*e*, where *e* is the electronic charge unit) are separated by the chromophore length (*l* = 7Å), such that μ_*eg*_ = |*q*|*l* = ∼12.8 D. In this case, the length *l* = 7Å was chosen to match the dimension of the trimethine group of the iCy3 molecule, as in prior work [14, 15, 36]. In applying the TQ model, the atomic coordinates *r*_*i*_ and transition charges *q*_*i*_ were determined from quantum chemical calculations [27]. In the following calculations, the atomic coordinates were held constant, and the transition charges were scaled uniformly by the factor 1.04319 to satisfy the condition μ_*eg*_ = |∑_*i*_ *q*_*i*_*r*_*i*_| = ∼12.8 D.

For all three models (PD, ED and TQ), the electrostatic coupling depends on the tilt angle, *θ*_*AB*_, the twist angle, *ϕ*_*AB*_, and the inter-chromophore separation, *R*_*AB*_ (Figs. 3*A* and 3*B*). However, for the atomistically-detailed TQ model the value of *J* also depends on the roll angle, *η*_*AB*_ (Fig. 3*C*). The sensitivity of the TQ model to changes in *η*_*AB*_, applied either symmetrically or anti-symmetrically, permits us to distinguish between varying degrees of stacking between the planar faces of the iCy3 trimethine groups. For completeness, we also allow for shear and shift displacements, *δ*_*AB*_ and *ξ*_*AB*_, respectively (Figs. 3*D* and 3*E*).

In Fig. 4 are shown examples of (iCy3)_2_ dimer conformations in which individual atoms have been color-coded according to their transition charge (TQ) values. We adopt the convention that *η*_*AB*_ = 0 corresponds to the ‘edge-to-edge’ conformation in which the linkages that tether the iCy3 probe to the DNA sugar-phosphate backbone are positioned on opposite sides of the dimer (see Fig. 4*B*). Starting from the edge-to-edge conformation, the anti (syn) face-to-face conformation is realized by rotating symmetrically (antisymmetrically) about the *η*_*AB*_ axes, as shown in Fig. 4*A* (Fig. 4*C*). Thus, the coordinates *θ*_*AB*_, *ϕ*_*AB*_, *R*_*AB*_, *η*_*AB*_, *δ*_*AB*_ and *ξ*_*AB*_ can be used to fully specify the (iCy3)_2_ dimer conformation. We note that the inter-chromophore separation, shear and shift displacements are largely limited by steric constraints imposed by the DNA duplex framework, located either on both sides of the (iCy3)_2_ dimer probe (position +15) or on only one side of the dimer probe (positions +2, +1 and -1). The situation at position -2 is more ambiguous and clearly less constrained (see Fig. 1).

**Figure 4.**
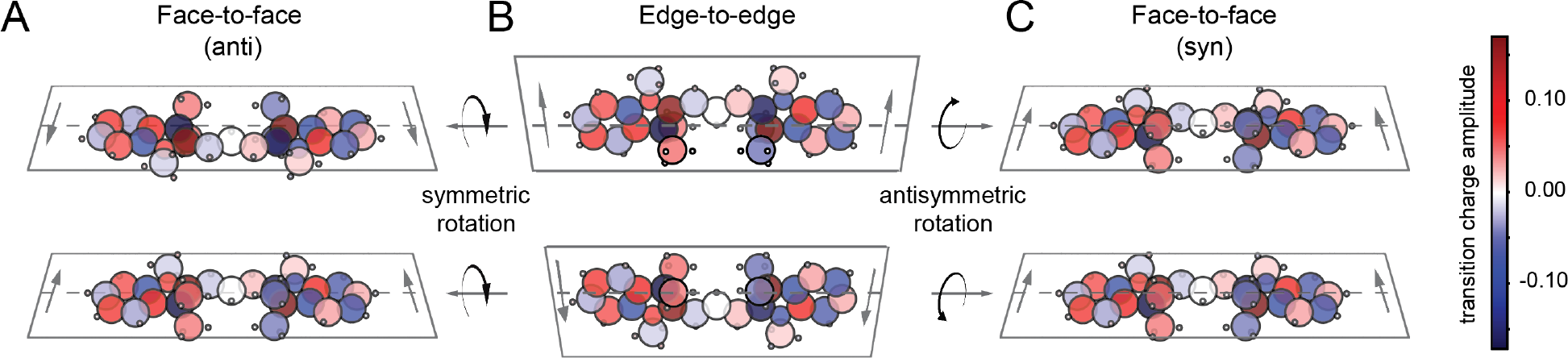
Inclined view of three-dimensional transition charge (TQ) density maps for the (iCy3)_2_ dimer with (*A*) face-to-face (anti) conformation, (*B*) edge-to-edge conformation, and (*C*) face-to-face (syn) conformation. The edge-to-edge conformation shown in *B* corresponds to values of the conformational parameters: *ϕ*_*AB*_ = *θ*_*AB*_ = *η*_*AB*_ = *δ*_*AB*_ = *ξ*_*AB*_ = 0 and *R*_*AB*_ = 12.8Å. Gray arrows point toward the side of the iCy3 chromophore probe that is directly bonded to the DNA sugar-phosphate backbone. The anti and syn face-to-face conformations (with *η*_*AB*_ = 90°) are achieved by rotating the roll angle symmetrically and antisymmetrically, respectively, as indicated.

### C. Numerical Optimization Procedure

In prior work [13-15], we characterized the iCy3 monomer absorbance spectrum using the effective-mode vibronic Hamiltonian [Eq. (1)] with parameters: ε_*eg*_ ≅ 18,250 cm^-1^, λ^2^ ≅ 0.57, ω_0_ ≅ 1,100 cm^-1^ and inhomogeneity parameter σ_*A*_ ≅ 300 cm^-1^. We determined these values by performing a numerical optimization procedure in which we compared the simulated monomer absorbance spectrum to experimental data. We assumed constant values for the homogeneous FWHM linewidth Γ_*H*_ = 186 cm^-1^ and the monomer EDTM μ_*eg*_ = ∼12.8 D, which we determined in separate experiments [13]. In the current work, we used the above parameters as inputs to our analyses of the absorbance and CD spectra of the (iCy3)_2_ dimer-labeled ss-dsDNA constructs [see Eqs. (5) and (6), respectively]. The experimental dimer absorbance and CD spectra were thus used to determine optimized values of the structural parameters *R*_*AB*_, *ϕ*_*AB*_ and *θ*_*AB*_, which specified the electrostatic coupling *J* under the PD and ED models [14, 15]. As in our prior studies, we used 6 vibrational states in the ground and excited electronic states for each monomer (12 total) giving rise to a dimer product state space of dimension 12 × 12 = 144.

In the current work we performed calculations to efficiently explore a seven-dimensional parameter space to obtain optimized values of the structural coordinates that characterize the (iCy3)_2_ dimer conformation. These coordinates are: *i*) the three angular transformations *η*_*AB*_, *θ*_*AB*_ and *ϕ*_*AB*_; *ii*) the three linear spatial translations *R*_*AB*_, *δ*_*AB*_ and *ξ*_*AB*_; and *iii*) the inhomogeneous lineshape parameter σ_*A*_. In our procedure, we carried out a systematic grid search in combination with a differential evolution algorithm to identify candidate trial conformations [28, 29]. We obtained each trial conformation by performing an ordered sequence of coordinate transformations on a dimer initially in an ‘edge-to-edge’ configuration, as shown in Fig. 4*B*, with *ϕ*_*AB*_ = *θ*_*AB*_ = *η*_*AB*_ = *δ*_*AB*_ = *ξ*_*AB*_ = *R*_*AB*_ = 0. For the starting configuration, the inter-chromophore spacing, *R*_*AB*_, was set to zero such that the two iCy3 monomers superimposed with their long axes aligned with the *z*-axis. The sequence of operations was: (*i*) Each monomer was rotated about the *z*-axis by the ‘roll’ angle *η*_*AB*_ (see Fig. 3*C*). For symmetric rotations both monomer *A* and monomer *B* were rotated by *η*_*AB*_, and for antisymmetric rotations monomer *A* was rotated by *η*_*AB*_ and monomer *B* by −*η*_*AB*_ (Fig. 4). (*ii*) The relative ‘tilt’ angle was applied (Fig. 3*A*) by rotating monomer *A* and monomer *B* about the *y*-axis by *θ*_*AB*_/2 and −*θ*_*AB*_/2, respectively. (*iii*) The ‘twist’ angle was applied (Fig. 3*B*) by rotating monomer *A* and monomer *B* about the *x*-axis by *ϕ*_*AB*_/2 and −*ϕ*_*AB*_/2, respectively. (*iv*) The ‘shear’ displacement was applied (Fig. 3*D*) by translating monomer *A* and monomer *B* along the *z*-axis by *δ*_*AB*_/2 and −*δ*_*AB*_/2, respectively. (*v*) The ‘shift’ displacement was applied (Fig. 3*E*) by translating monomer *A* and monomer *B* along the *y*-axis by *ξ*_*AB*_/2 and −*ξ*_*AB*_/2, respectively. (*vi*) The inter-chromophore separation was applied (Fig. 3*B*) by translating monomer *A* relative to monomer *B* along the *x*-axis by 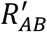. The final center-to-center separation was determined using the formula 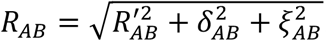.

For each dimer conformation, we calculated a linear least squares error function χ^2^, which we subsequently minimized using the SciPy differential evolution and minimization algorithms in the Python programming platform [38].

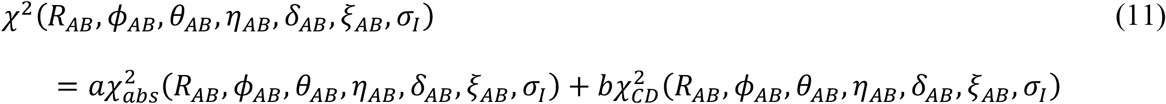

In Eq. (11), the coefficients *a* and *b* were chosen so that the relative contributions from absorbance and CD spectra were equally weighted. We performed error-bar analyses of the optimized parameters, which we determined by a 1% deviation of the χ, function from its optimized value. We additionally limited the dimension of the search by imposing constraints on the three linear spatial translations *R*_*AB*_, *δ*_*AB*_ and *ξ*_*AB*_. We limited the separation distance to fall within the range 5Å ≤ *R*_*AB*_ ≤ 10Å. We also constrained the shear and shift displacements to fall within the ranges: -3Å ≤ *δ*_*AB*_ ≤ +2Å, -2Å ≤ *ξ*_*AB*_ ≤ +3Å for the +15 and +2 constructs; -3Å ≤ *δ*_*AB*_, *ξ*_*AB*_ ≤ +3Å for the +1 constructs, and -4Å ≤ *δ*_*AB*_, *ξ*_*AB*_ ≤ +4Å for the -1 and -2 constructs. We found that computational efficiency was significantly improved by implementing the differential evolution algorithm [28, 29] to generate trial conformations. A candidate structure that violated the van der Waals overlap criteria (*r*_*w*_ = 1.5Å) was rejected by returning to the value of χ^2^ associated with the simulated iCy3 monomer spectra. Such steric violations led to discontinuities in the χ^2^ surface, which we circumvented by implementing a stochastic sampling approach, as in [10]. For each of the three electrostatic coupling models that we considered (PD, ED and TQ), we included the effects of steric interactions. This was accomplished by performing the same three-dimensional orientational coordinate transformations of the (iCy3)_2_ dimer during the search algorithm while accounting for the van der Waals radii of the iCy3 monomers. We note that while the PD and ED models are insensitive to the roll angle *η*_*AB*_, the simple inclusion of short-range steric interactions in these models provides ostensibly the additional information needed to determine this parameter.

Finally, our choice for the generalized van der Waals radius *r*_*w*_ = 1.5Å represents a reasonable average value given accepted values for H (1.1Å), C (1.8Å), N (1.6Å) and O (1.6Å) [39]. We found that using slightly larger or smaller values for *r*_*w*_ did not lead to significantly different results.

### D. Transition Electrostatic Potential (trESP) Charges for TQ Model Calculations

A comprehensive description of the computational procedure used to obtain the transition electrostatic potential (trESP) charges for the Cy3 molecule was reported previously [27]. Briefly, the ground electronic state of the Cy3 monomer was optimized in an implicit solvent (water) using B3LYP/6-31G(d). Using the optimized ground state configuration, vertical excited electronic state calculations were performed using LC-ωPBE/Def2SVP in an implicit solvent (water). The Multiwfn (A Multifunctional Wavefunction Analyzer) program [40] was used to determine the transition density matrix between the ground and first excited electronic state. Using the transition density matrix, the transition electrostatic potential was calculated on a grid surrounding the molecule, then the trESP was decomposed into transition point charges centered on the Cy3 atomic coordinates using the CHELPG (Charges from Electrostatic Potentials using a Grid-based method) fitting procedure [41]. The resulting trESP charges reproduced the magnitude and orientation of the Cy3 monomer EDTM determined by direct *ab initio* calculations using LC-ωPBE/Def2SVP.

### E. Sample Preparation

The sequences and nomenclature of the (iCy3)_2_ dimer-labeled ss-dsDNA constructs used in the current work are the same as those used in previous studies [15]. Oligonucleotide samples were purchased from Integrated DNA Technologies (IDT, Coralville, IA) and used as received. Solutions were prepared using ∼1 μM oligonucleotide in 10 mM TRIS buffer (pH ∼ 7) with 100 mM NaCl and 6 mM MgCl_2_. Complementary strands were combined in equimolar concentrations. The samples were heated to 95°C for 4 min and left to cool slowly on a heat block overnight prior to data collection. The annealed (iCy3)_2_ dimer-labeled ss-dsDNA fork constructs contained both ss and ds DNA regions, with the probe labeling positions indicated by the nomenclature described in Fig. 1*C*.

### F. Absorbance and Circular Dichroism (CD) Measurements

We performed absorbance measurements using a Cary 3E UV-vis spectrophotometer and CD measurements using a JASCO model J-720 CD spectrophotometer. All absorbance and CD measurements were performed at room temperature, and the samples were housed in a 1 cm quartz cuvette.

### G. Visualization Models of (iCy3)_2_ Dimer-Labeled ss-dsDNA Fork Constructs

To examine how the (iCy3)_2_ dimer conformations, which we determined from our spectral analyses, fit into their corresponding DNA frameworks, we constructed molecular models using the PyMOL [42] and Avogadro [43] software. In building these models, we assumed that: (*i*) the atomic coordinates of the component iCy3 monomers are those we previously determined in quantum chemical calculations [27]; and (*ii*) that the DNA bases and sugar-phosphate backbones immediately adjacent to the (iCy3)_2_ dimer probes are not greatly perturbed by their presence.

For a given (iCy3)_2_ dimer conformation, we generated a protein data bank (PDB) file containing the *xyz* atom coordinates. We next used the Builder / Nucleic Acid function of PyMOL to create B-form double-stranded DNA segments and single-stranded DNA segments of the base-sequence-specific (iCy3)_2_ dimer-labeled ss-dsDNA constructs that we studied [15]. We used the Builder / Chemical Tool function of PyMOL to create the ethoxy linkers (-CH_2_-CH_2_-O-), which we used to connect the carbon of the iCy3 to the phosphorous of the phosphate group in the DNA backbone, as shown in Fig. 1*A*. We note that for a given (iCy3)_2_ dimer conformation, the relative positions of the DNA segments that are conjoined by the dimer are significantly constrained. In a final step, we used the Avogadro software to optimize the geometries of the 8 carbons within the 4 ethoxy linkers while fixing the atomic coordinates of the (iCy3)_2_ dimer. This resulted in only very small changes to the coordinates of the ethoxy linkers and the neighboring bases. From the *xyz* coordinates of the deoxyribose C1′ atom connected to each of the four bases, we determined the ‘helical rotation’ and the ‘helical spacing’ between the DNA segments introduced by the insertion of the dimer probe, as shown in Fig. 5.

**Figure 5.**
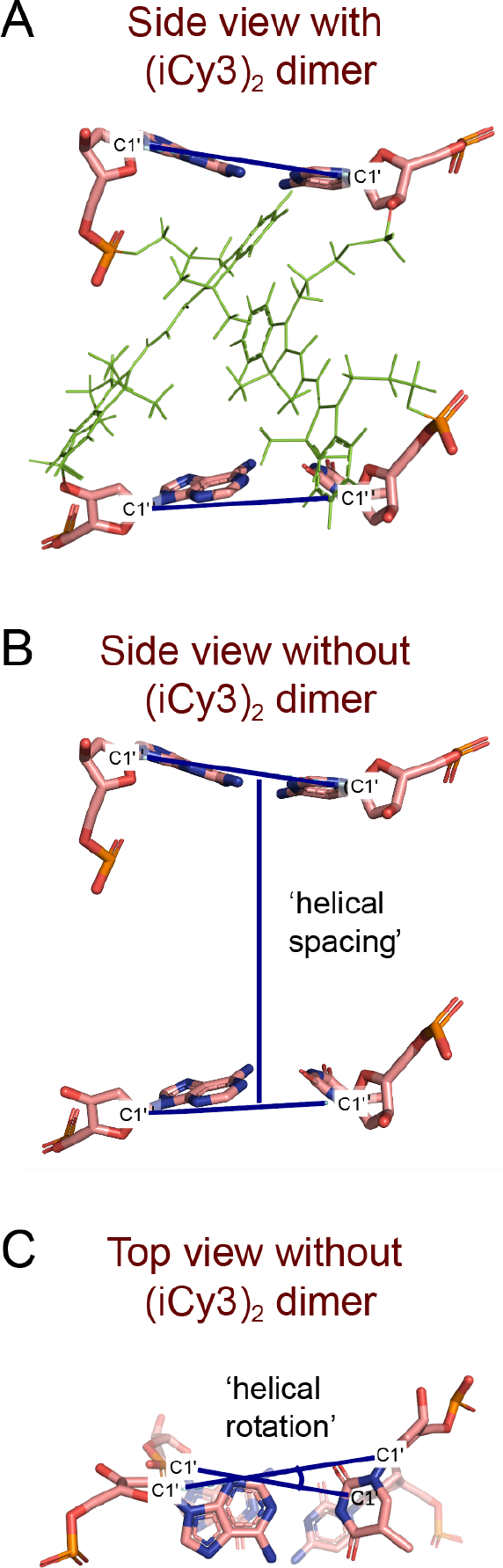
PyMOL model of (iCy3)_2_ dimer-labeled ss-dsDNA fork construct showing the four nucleotides immediately adjacent to the (iCy3)_2_ dimer probe. (*A*) Line segments are shown (dark blue) connecting the deoxyribose C1′ atoms of the two base pairs ‘above’ and ‘below’ the (iCy3)_2_ dimer probe. (*B*) The ‘helical spacing’ is defined as the distance between the midpoints of the line segments connecting the C1′ atoms of the upper and lower base pairs. (*C*) The ‘helical rotation’ is defined as the angle between the projections of these line segments onto the transverse plane.

**Figure 6.**
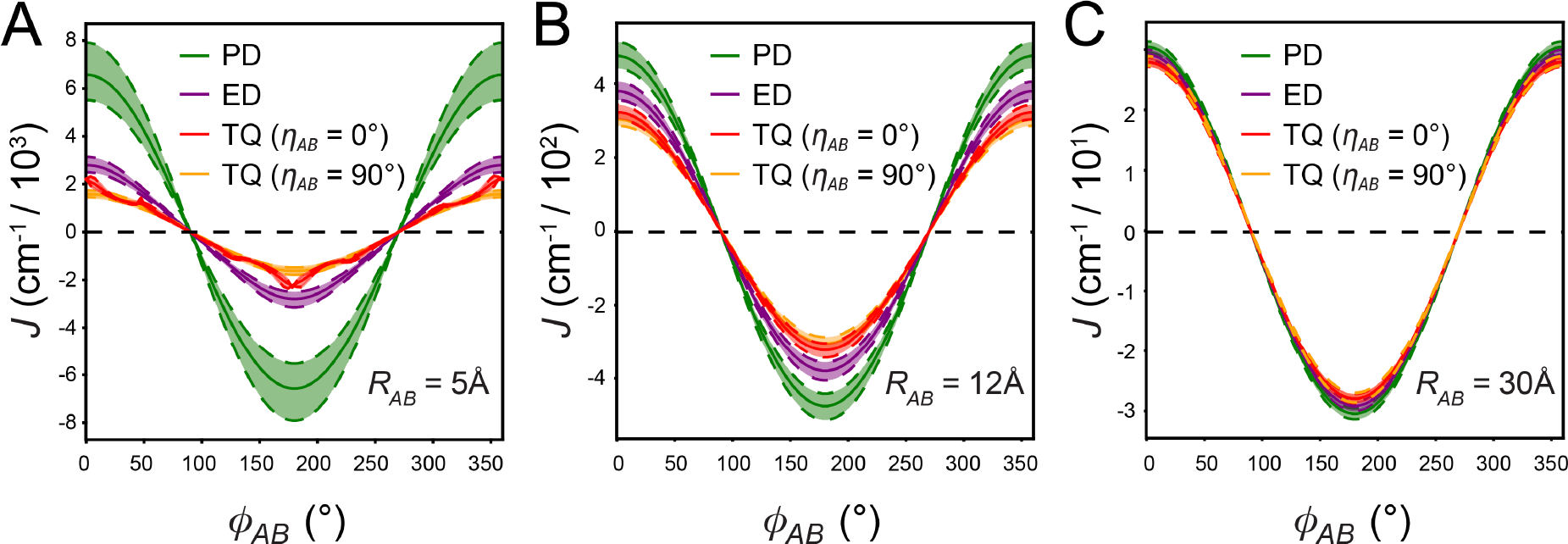
Dependence of the electrostatic coupling *J* on the twist angle *ϕ*_*AB*_ under the PD, ED, and TQ models for (*A*) *R*_*AB*_ = 5Å, (*B*) *R*_*AB*_ = 12Å and (*C*) *R*_*AB*_ = 20Å. The values for *θ*_*AB*_ = *δ*_*AB*_ = *ξ*_*AB*_ = 0. For the TQ calculations, both the ‘edge-to-edge’ (*η*_*AB*_ = 0) and (anti) ‘face-to-face’ conformations (*η*_*AB*_ = 90°) are shown. Results for the (anti) and (syn) ‘face-to-face’ conformations (with *η*_*AB*_ = 90°) are indistinguishable at all values of *R*_*AB*_.

## III. Results

### A. Transition Density Model Calculations for the (iCy3)_2_ Dimer

In Fig. 6, we present calculations of the electrostatic coupling *J* for the PD, ED and TQ models. Here we set *θ*_*AB*_ = *δ*_*AB*_ = *ξ*_*AB*_ = 0 and plot *J* as a function of the twist angle *ϕ*_*AB*_ for three different intermolecular separations, *R*_*AB*_ = 5, 12 and 30Å. For the TQ calculations, we examine specifically the ‘edge-to-edge’ (*η*_*AB*_ = 0) and anti ‘face-to-face’ (*η*_*AB*_ = 90°) conformations. Here we ignore steric overlap between molecules to focus on the Coulomb interactions between effective point charges. We plot only the results for the anti ‘face-to-face’ conformation because the results plotted at the given resolution are indistinguishable from those of the syn conformation. In all cases, *J* varies sinusoidally between maximum and minimum values at *ϕ*_*AB*_ = 0° and 180°, respectively, and undergoes zero-crossings at *ϕ*_*AB*_ = 90° and 270°. At the smallest separation, *R*_*AB*_ = 5Å, which is significantly less than the iCy3 molecular dimension (∼14Å), the coupling strength for all three models is generally large and sensitive to *ϕ*_*AB*_ (Fig. 6*A*). Moreover, at such small separations the three coupling models exhibit the greatest variability. Of the three models, TQ represents the most accurate approximation of the transition density, which assigns trESP transition charge values to the individual atoms of the iCy3 molecule [27]. We see that the PD model overestimates the coupling strength due to its single point-dipole at the center-of-mass position of the iCy3 probe. Similarly, the ED model overestimates the coupling strength, albeit to a far lesser extent than the PD model. For the ‘edge-to-edge’ conformation of the TQ model (*η*_*AB*_ = 0) there is an additional low amplitude *ϕ*_*AB*_-dependent modulation that is due to the Coulomb interactions between non-physically overlapping point charges, which does not occur for the ‘face-to-face’ configuration (*η*_*AB*_ = 90°). The similar magnitudes of *J* predicted by the ED and TQ models suggests that by simply accounting for the finite dimension of the molecule it is possible to determine a reasonable estimate of the *ϕ*_*AB*_-dependent coupling, as suggested previously [35, 36].

As *R*_*AB*_ is increased to values close to and exceeding the molecular dimension, the coupling strength decreases rapidly, as does the variability between the three models (see Figs. 6*B* and 6*C*). For *R*_*AB*_ = 12Å, the magnitude of *J* is on the order of a few hundred wavenumbers. We note that the experimental value of *J* for (iCy3)_2_ dimer-labeled dsDNA constructs is on the order of ∼500 cm^-1^ [13-15]. Thus, these results indicate that *R*_*AB*_ must be smaller than the cross-sectional diameter of the Watson-Crick B-helix (∼ 20Å), which is as expected. We note that for the largest separation shown, *R*_*AB*_ = 30Å, the magnitude of *J* is on the order of a few tens-of-wavenumbers (see the relative magnitudes of the vertical axes in Fig. 6) so that the results using the three models are nearly indistinguishable.

To rule out sterically overlapping conformations from our optimization procedure we adopted a generalized van der Waals radius *r*_*w*_ = 1.5Å, which we applied uniformly to the constituent atoms. When excluded volume is considered, the parameter space of the (iCy3)_2_ dimer is sterically constrained so that only non-overlapping conformations are included. In Fig. 7 are shown two-dimensional heat maps of the electrostatic coupling *J* plotted as a bivariate function of the twist angle *ϕ*_*AB*_ and the roll angle *η*_*AB*_. The top row of Fig. 7 (panels *A* – *D*) shows the *η*_*AB*_-dependence of *J* upon symmetric rotation, and the bottom row (panels *E* – *H*) shows the *η*_*AB*_-dependence of *J* upon anti-symmetric rotation. The two sets of calculations (symmetric and antisymmetric) exhibit qualitatively very similar behavior, although there are quantitative differences.

**Figure 7.**
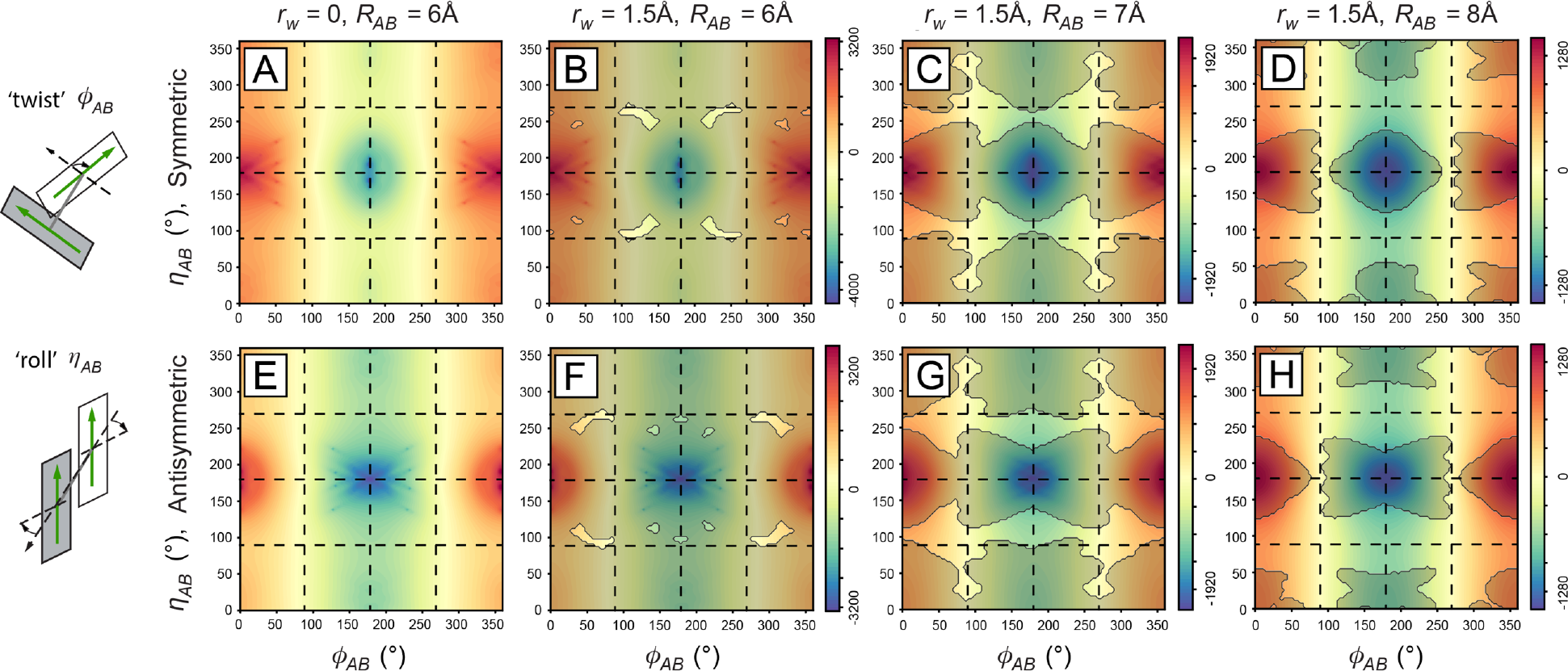
The electrostatic coupling *J* under the TQ model is plotted as a function of the twist angle *ϕ*_*AB*_ and the roll angle *η*_*AB*_ (schematically illustrated on the left) for *δ*_*AB*_ = *ξ*_*AB*_ = 0. The top row shows the roll angle varied symmetrically and the bottom row antisymmetrically. The first column (*A, E*) shows the *ϕ*_*AB*_, *η*_*AB*_-dependence of *J* without accounting for van der Waals overlap (*r*_*w*_ = 0). The remaining columns consider the effects of van der Waals overlap with *r*_#_ = 1.5Å for *R*_*AB*_ = 6Å (*B, F*), *R*_*AB*_ = 7Å (*C, G*), and *R*_*AB*_ = 8Å (*D, H*). Gray semi-transparent masks indicate sterically-disallowed regions of the conformational space. Vertical and horizontal dashed lines indicate the angles 90°, 180° and 270°. Color bars indicate the value of *J* in units of cm^-1^.

The first column of Fig. 7 (panels *A, E*) shows the *η*_*AB*_, *ϕ*_*AB*_-dependence of *J* for *R*_*AB*_ = 5Å, while ignoring the effects of steric interactions (*r*_*w*_ = 0). We see that the *ϕ*_*AB*_-dependent modulation of *J* is a parametric function of the roll angle *η*_*AB*_. For the ‘edge-to-edge’ conformation (*η*_*AB*_ = 0), the *ϕ*_*AB*_-dependence of *J* is the same as that shown in Fig. 6. The modulation amplitude is slightly enhanced for the inverted ‘edge-to-edge’ conformation with *η*_*AB*_ = 180° relative to *η*_*AB*_ = 0. However, for the ‘face-to-face’ conformations (with *η*_*AB*_ = 90° and 270°), the modulation amplitude is significantly reduced.

The remaining panels of Fig. 7 show the effects of steric interactions between the iCy3 chromophores at close separation by indicating regions of the conformational space that are excluded due to van der Waals overlap. Each successive column in Fig. 7, from left to right, corresponds to an increasing value of the inter-chromophore separation *R*_*AB*_ = 6Å (Fig. 7*B* and 7*F*), 7Å (Fig. 7*C* and 7*G*) and 8Å (Fig. 7*D* and 7*H*). Here, sterically-disallowed conformations are indicated by the masked semi-transparent gray regions, and sterically-allowed conformations by the unmasked regions. For the smallest separation *R*_*AB*_ = 6Å, nearly all the conformational space is sterically disallowed. However, for *R*_*AB*_ = 7Å a significant fraction of the conformational space becomes accessible, which favors primarily the two ‘face-to-face’ conformations (*η*_*AB*_ = 90° and 270°) relative to the two ‘edge-to-edge’ conformations (*η*_*AB*_ = 0° and 180°). These calculations also indicate that steric interactions favor orthogonal values for the twist angle (*ϕ*_*AB*_ = 90° and 270°) relative to parallel and antiparallel conformations (*ϕ*_*AB*_ = 0° and 180°, respectively).

The calculations summarized in Fig. 7 permit us to generate (iCy3)_2_ dimer conformations that avoid steric overlap. These results suggest minor refinements to some previously optimized conformations of (iCy3)_2_ dimer-labeled ss-dsDNA fork constructs using the PD and ED models for which we reported values of *R*_*AB*_ < 5Å [14, 15]. The refined structures reported in the current work account for sterically disallowed conformations at near contact separations, which is important in terms of their reflections of the adjacent DNA structures. Another important consideration is that the space of sterically-allowed (iCy3)_2_ dimer conformations exhibits a relatively high level of *J*-degeneracy. Thus, the conformation of an (iCy3)_2_ dimer-labeled DNA construct cannot be determined from the value of the coupling strength alone, and additional constraints must be included to obtain a unique set of conformational parameters that are consistent with experimental data. These constraints are provided by the electric dipole transition moments (EDTMs) and the rotational strengths of the coupled dimer, which determine the absorbance and CD spectral line shapes, respectively, as described by Eqs. (3) – (6).

We next consider the dependence of the electrostatic coupling *J* on the shear and shift displacements (*δ*_*AB*_ and *ξ*_*AB*_, respectively, see Fig. 3*D* and 3*E*) under the TQ model. In Fig. 8 are shown the *δ*_*AB*_, *ξ*_*AB*_-dependence of *J* for the anti ‘face-to-face’ conformation (*η*_*AB*_ = 90°, *ϕ*_*AB*_ = *θ*_*AB*_ = 0). In Fig. 8*A*, the separation *R*_*AB*_ = 8Å is large enough to avoid steric overlaps for all values of *δ*_*AB*_ and *ξ*_*AB*_. *J* has its peak value at *δ*_*AB*_ = *ξ*_*AB*_ = 0 and falls off anisotropically for finite displacements. For *δ*_*AB*_ = ∼9Å, the coupling *J* changes sign as partial transition charges of the opposite (same) sign move closer to (away from) one another. As the inter-chromophore separation is decreased to *R*_*AB*_ = 7Å, 6Å and 5Å (see Figs. 8*B* – 8*D*, respectively), steric interactions become increasingly significant. The effect of steric interactions at close separation is to exclude some high symmetry conformations of the (iCy3)_2_ dimer (e.g., with *δ*_*AB*_ ≈ *ξ*_*AB*_ ≈ 0) and to concomitantly reduce the coupling strength. For the (iCy3)_2_ dimer-labeled ss-dsDNA constructs, variations of the shift and shear parameters are expected to be highly constrained by the cylindrical symmetry imposed by local complementary (Watson-Crick) bases and sugar-phosphate backbones. Thus, for close separations, very small adjustments of the shear and shift displacements can reduce steric overlap and produce model absorbance and CD spectra that compare favorably with the experimental data, which is consistent with studies by Hestand and Spano [44].

**Figure 8.**
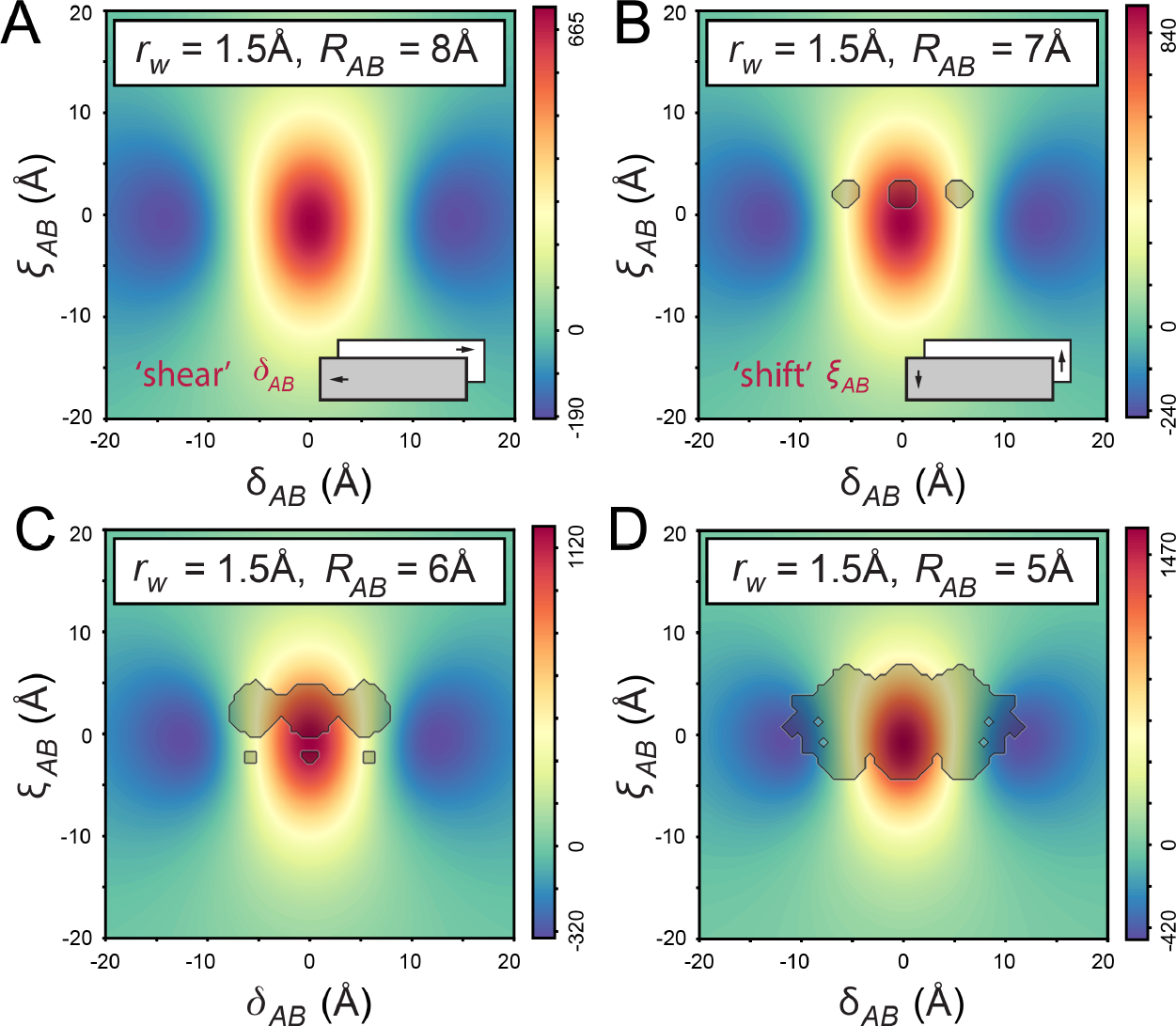
The electrostatic coupling, *J*, under the TQ model is plotted as a function of the ‘shear’ (*δ*_*AB*_) and ‘shift’ (*ξ*_*AB*_) displacements for the anti ‘face-to-face’ conformation (*η*_*AB*_ = 90°, *ϕ*_*AB*_ = *θ*_*AB*_ = 0). The shear and shift parameters are illustrated schematically in the insets of panels *A* and *B*, respectively. The effects of steric interactions are shown as a function of the separation with *r*_*w*_ = 1.5Å and (*A*) *R*_*AB*_ = 8Å, (*B*) *R*_*AB*_ = 7Å, (*C*) *R*_*AB*_ = 6Å and (*D*) *R*_*AB*_ = 5Å. Gray semi-transparent masks indicate sterically disallowed regions of the conformational space. Color bars indicate the value of *J* in units of cm^-1^.

### B. Parameterization of Absorbance and CD Spectra of +15 (iCy3)_2_ Dimer-Labeled ss-dsDNA Constructs

In Fig. 9 are shown the results of our optimization procedure for the TQ model applied to the absorbance and CD spectra of the +15 (iCy3)_2_ dimer-labeled ss-dsDNA construct. In each of the 9 panels, the optimized χ_2_ values are plotted as a joint function of the twist angle *ϕ*_*AB*_ and the tilt angle *θ*_*AB*_ for fixed *R*_*AB*_ (incremented by row, 5.8Å, 6.7Å and 7.6Å) and fixed roll angle *η*_*AB*_ (incremented by column, 90°, 110° and 130°). In these calculations, the shear and shift parameters were set to the optimized values *δ*_*AB*_ = ∼0.2Å and *ξ*_*AB*_ = ∼3.0 Å, which we determined by extensively searching the parameter space. Green-shaded regions indicate the most favorable solutions to the optimization procedure, while gray semi-transparent masks indicate regions of the conformational space that are sterically disallowed (with *r*_*w*_ = 1.5Å).

**Figure 9.**
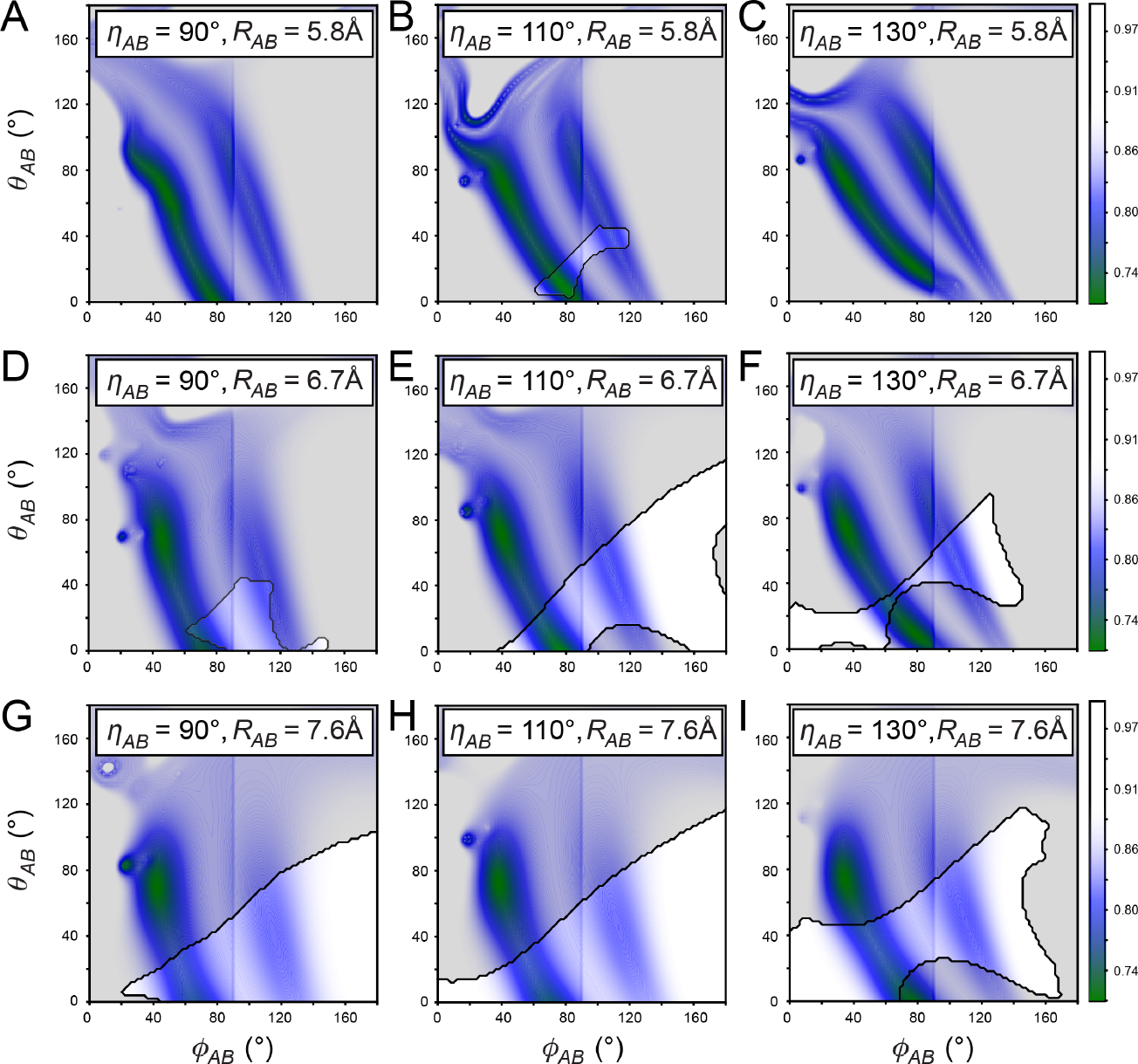
The χ^2^ error function is determined from fitting the TQ model to experimental absorbance and CD data for the +15 (iCy3)_2_ dimer-labeled ss-dsDNA construct. Each panel shows the logarithm of χ^2^, normalized to the highest value of the surface, and plotted as a function of *ϕ*_*AB*_ and *θ*_*AB*_. Successive rows (labeled *A, D, G*) indicate increasing values of *R*_*AB*_ (5.8Å, 6.7Å and 7.6Å, respectively), and successive columns (labeled *A, B, C*) indicate increasing values of *η*_*AB*_ (varied symmetrically from 90°, 110° and 130°). For these calculations, we have set *δ*_*AB*_ = ∼ 0.2Å and *ξ*_*AB*_ = ∼3.0Å. Semi-transparent gray masks indicate regions of the conformational space that are inaccessible due to steric interactions (*r*_*w*_ = 1.5Å). Green-shaded regions indicate best agreement between the TQ model and experimental data.

The calculations presented in Fig. 9 show that the TQ model parameter space for the +15 (iCy3)_2_ dimer-labeled DNA construct contains a locus of optimal solutions (shaded green, with twist angle 45° ≲ *ϕ*_*AB*_ ≲ 90°), which occur over a relatively broad range of values for *θ*_*AB*_, *η*_*AB*_ and *R*_*AB*_. In the bottom half of each panel (for *θ*_*AB*_ ≲ 80°) there are two channels of quasi-optimal solutions: a primary channel (shaded green) with right-handed conformations (*ϕ*_*AB*_ < 90°) and a secondary channel (shaded blue) with left-handed conformations (*ϕ*_*AB*_ > 90°). While both channels agree favorably with the experimental absorbance spectrum, only the right-handed conformations agree with the experimental CD spectrum. The parameter space is significantly restricted at small separations due to steric interactions between the iCy3 monomers (gray-shaded regions). The fraction of accessible conformations increases abruptly as the separation is increased within the range 5.8Å ≲ *R*_*AB*_ ≲ 6.7Å (compare successive rows), and for values of the roll angle within the range 90° ≲ *η*_*AB*_ ≲ 130° (compare middle column to left and right columns). For values of *R*_*AB*_ > ∼7Å (bottom row), the optimal solutions occur at relatively large tilt angles that correspond to one or more steric conflicts. These and similar calculations indicate that only a relatively narrow range of structural parameters can simultaneously provide good agreement with the experimental data and avoid steric overlap. By systematically searching the parameter space, we obtained a globally optimized solution with *ϕ*_*AB*_ = ∼80°, *θ*_*AB*_ = ∼2.4°, *η*_*AB*_ = ∼105°, *δ*_*AB*_ = ∼ 0.2Å, *ξ*_*AB*_ = ∼3.0Å and *R*_*AB*_ = ∼5.9Å. The corresponding *ϕ*_*AB*_, *θ*_*AB*_-dependent χ^2^ surface is shown in Fig 10*A*, and the global minimum is indicated by the red dot. A schematic 3D model of the optimized (iCy3)_2_ dimer conformation is shown in Fig 10*B*.

**Figure 10.**
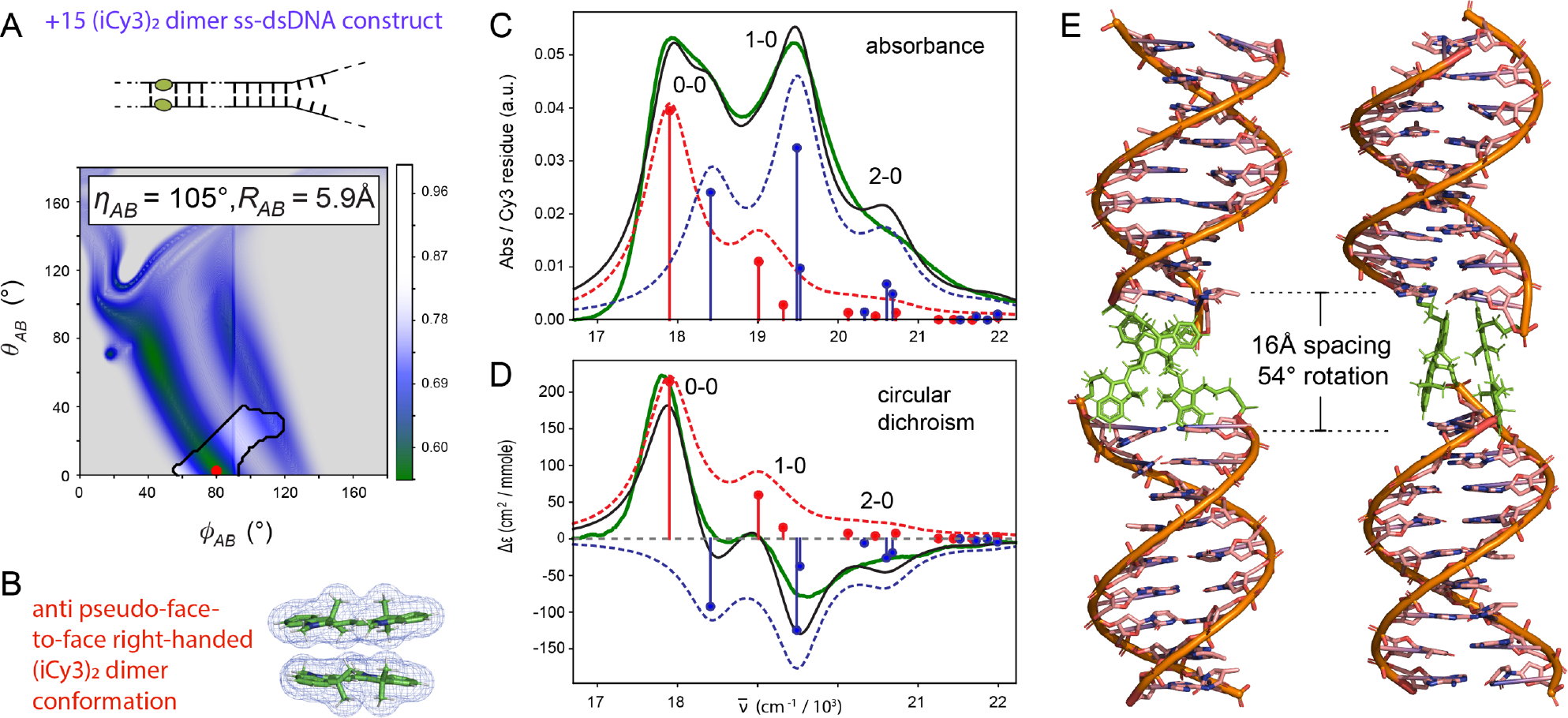
(*A*) χ^2^ error function determined from fitting the TQ model to experimental absorbance and CD data for the anti +15 (iCy3)_2_ dimer-labeled ss-dsDNA construct plotted as a function of *ϕ*_*AB*_ and *θ*_*AB*_, and for *η*_*AB*_ = 105.4°, *δ*_*AB*_ = ∼ 0.2Å, *ξ*_*AB*_ = ∼3.0Å and *R*_*AB*_ = ∼5.9Å. Semi-transparent gray regions indicate sterically disallowed conformations using an average atomic van der Waals radius *r*_*w*#_ = 1.5Å. The red dot indicates the globally minimized value of χ^2^, which occurs within the sterically allowed region. (*B*) Side view of the optimized anti (iCy3)_2_ dimer conformer. Comparison between simulated and experimental (*C*) absorbance and (*D*) CD spectra. Spectra were recorded using 100 mM NaCl and 6 mM MgCl_2_ in 10 mM TRIS buffer (pH ∼7) at 25°C. Simulated spectra correspond to the optimized values of the parameters (see Table 1). Experimental absorbance and CD spectra are shown (in green) overlaid with theoretical spectra (black curves) spanning the 0-0, 1-0 and 2-0 vibronic sub-bands. Symmetric (+) and anti-symmetric (−) excitons are shown as blue and red sticks (and dashed blue and red curves), respectively. (*E*) PyMOL visualizations of the optimized anti (iCy3)_2_ dimer conformation built into the framework of the DNA duplex, which was geometrically optimized by fixing coordinates of the (iCy3)_2_ dimer atoms as described in Sect. II.G. The two models are shown rotated 90° relative to one another.

Direct comparisons between the optimized absorbance and CD spectra based on the TQ model and experimental data for the +15 (iCy3)_2_ dimer-labeled ss-dsDNA construct are shown in Figs. 10*C* and 10*D*, respectively. In general, the agreement between experiment and theory using the combined Holstein-Frenkel (H-F) Hamiltonian and TQ model of the transition density is very good. Both simulated absorbance and CD spectra reflect the experimentally-observed effects of intensity borrowing between the 0-0 and 1-0 sub-bands – i.e., the diminished oscillator strength of the lowest-energy (0-0) antisymmetric transition and the enhanced oscillator strength of the vibrationally excited (1-0) symmetric transition [13, 25]. However, the theory generally fails to predict the exact spectral line shapes, as it overestimates the intensities of the blue-edge shoulder of the 0-0 feature and the vibrationally excited 1-0 and 2-0 features. This is likely due to inaccuracies of the vibronic Hamiltonian used to describe the iCy3 monomer, which assumes a single effective harmonic mode [21]. Nevertheless, the H-F model predicts that the essential features of the absorbance and CD spectra of the +15 (iCy3)_2_ dimer-labeled ss-dsDNA construct are consistent with experimental results over a broad range of environmental parameters [13-15].

In Fig. 10*E* are shown model visualizations of the optimized anti-face-to-face (iCy3)_2_ dimer conformation built into the DNA framework of the +15 DNA construct. This and the models presented below for the +2, +1 and -1 (iCy3)_2_ dimer-labeled ss-dsDNA fork constructs assume that the surrounding DNA bases and backbones, which are attached to the iCy3 probes by three carbon linkers (see Fig. 1*A*), are not significantly altered in secondary structure, apart from these probe attachments. Following this assumption, we determined the inter-base-pair separation and helical rotation of base pairs above and below the insertion sites of the (iCy3)_2_ dimer probes using line segments connecting the C1′ carbons of the glycosidic bonds between complementary bases. The dimer probe in the +15 DNA construct thus introduces a disruption to the normal spacing and helical rotation between consecutive base pairs. The distance between base pairs across the gap introduced by the (iCy3)_2_ dimer probe is ∼16Å, which is equivalent to the spacing of ∼4.4 stacked bases (with ∼3.6Å rise per base). However, the helical rotation introduced by the dimer probe is ∼54°, which is approximately the rotation equivalent to 1.6 consecutive base pairs in unlabeled B-DNA (with 34.3° rotation per residue) [45].

These structural considerations of the DNA framework relative to the (iCy3)_2_ dimer probe at the +15 position (i.e., probes positioned deep within the double-helical structure) may perturb the binding affinities of proteins that interact, either base-sequence specifically via hydrogen-bonding through the DNA grooves at positions immediately adjacent to the (iCy3)_2_ dimer probe, or non-specifically with the local sugar-phosphate backbones [46, 47]. However, the regular B-form geometry is likely reassumed at binding site positions located more than one or two base pairs removed from the probe-DNA interface (see Fig. 10*E*). Furthermore, breathing fluctuations at functional binding motifs for proteins located close to the probe-DNA interface may provide binding sites that, upon protein binding, are augmented by the binding interaction. Thus, differential effects on the structural parameters of the (iCy3)_2_ dimer probes in the absence and presence of specific binding proteins may also help to define structural aspects of biologically-relevant protein-DNA interactions.

We note that the parameters we determined for the +15 (iCy3)_2_ dimer-labeled ss-dsDNA construct differ somewhat from those we reported in previous studies, in which we determined the values *ϕ*_*AB*_ = ∼80°, *θ*_*AB*_ = ∼10° and *R*_*AB*_ = ∼4.4Å [15]. The previous work did not properly account for steric interactions, which led us to report sterically disallowed values less than 5Å for the inter-chromophore separation. We have found that discrepancies between the PD, ED and TQ models, which are most pronounced at very small separations, can lead to diverging results when steric interactions are ignored. Finally, accounting for steric interactions can provide information about the inter-dependences of conformational parameters. For example, as shown in Figs. 9*A* – 9*F*, at separation distances *R*_*AB*_ ≲ 7Å, steric interactions impose an upper bound on the possible values of *θ*_*AB*_ ≲ 40°. However, for larger values of *R*_*AB*_ ≳ 7Å (see Figs. 9*G* – 9*I*), the value of *θ*_*AB*_ must be increased to match the experimental spectra, in which case the value of *θ*_*AB*_ should not be restricted unnecessarily. This interdependence between *R*_*AB*_ and *θ*_*AB*_ occurs to some extent for all three of the electrostatic coupling models applied to the +15 (iCy3)_2_ dimer-labeled ss-dsDNA construct.

In Table 1, we present optimized solutions corresponding to each of the three electrostatic coupling models for the +15 (iCy3)_2_ dimer-labeled ss-dsDNA construct. We note that the optimized conformational parameters obtained using the three models (PD, ED and TQ for both symmetric and anti-symmetric roll) are very similar for the twist angle and differ somewhat for the tilt, roll and separation. The global minimum of the χ^2^ function corresponds to an anti-face-to-face conformation, with *ϕ*_*AB*_ = ∼80°, *θ*_*AB*_ = ∼2.4°, *η*_*AB*_ = ∼105°, *δ*_*AB*_ = ∼ 0.2Å, *ξ*_*AB*_ = ∼3.0Å and *R*_*AB*_ = ∼5.9Å, and a syn-face-to-face conformation with *ϕ*_*AB*_ = ∼79°, *θ*_*AB*_ = ∼12°, *η*_*AB*_ = ∼109°, *δ*_*AB*_ = ∼ -3.0Å, *ξ*_*AB*_ = ∼-0.5Å and *R*_*AB*_ = ∼6.7Å. We emphasize that these conformational assignments are only possible due to the detailed structural information provided by the TQ model.

**Table 1.** Optimized conformational parameters for the +15 (iCy3)_2_ dimer-labeled ss-dsDNA construct at 25°C based on the TQ, ED and PD models and van der Waals radius *r*_*w*_ = 1.5Å. Error bars were calculated based on a 1% deviation of the χ^2^ function from its minimum (optimized) value.

| | Model | $\phi_{AB}$ (°) | $\theta_{AB}$ (°) | $\eta_{AB}$ (°) | $\delta_{AB}$ (Å) | $\xi_{AB}$ (Å) | $R_{AB}$ (Å) | $\sigma$ (cm<sup>-1</sup>) | $J$ (cm<sup>-1</sup>) |
| --- | --- | --- | --- | --- | --- | --- | --- | --- | --- |
| Symm.<br>Roll,<br>'anti' | TQ | 80±1 | 2±1 | 105±2 | 0.2±0.8 | 3.0±0.4 | 5.9±0.1 | 320±17 | 515 |
|  | ED | 80±1 | 9±1 | 110±2 | 1.0±0.1 | 0.6±2.0 | 6.6±0.1 | 320±17 | 515 |
|  | PD | 79±1 | 14±1 | 123±40 | 0.0±0.5 | -0.7±0.1 | 7.6±0.1 | 320±16 | 515 |
| Antisym<br>Roll,<br>'syn' | TQ | 79 ±1 | 12±1 | 109±3.0 | -3.0±0.2 | -0.5±0.4 | 6.7±0.1 | 320±17 | 515 |
|  | ED | 80±1 | 8±16 | 89±15 | 0.6±1.0 | 0.8±0.1 | 6.3±0.1 | 321±17 | 514 |
|  | PD | 79±1 | 13±1 | 92±40 | -1.0±0.1 | 0.4±1.0 | 7.6±0.1 | 321±16 | 514 |

Although the optimized anti and syn conformations provide equivalent agreement with the experimental spectra, the anti-conformation provides the most reasonable accounting for steric interactions within the (iCy3)_2_ dimer probe. It is perhaps somewhat surprising that roughly similar values for all optimized parameters (including *η*_*AB*_) were obtained using the three electrostatic coupling models (TQ, PD and ED). This result was obtained only when steric interactions between the iCy3 monomers were accounted for. We note that the associated error bars for the roll angle, *η*_*AB*_, obtained using the PD and ED models are somewhat larger than those obtained with the TQ model.

### C. Parameterization of Absorbance and CD Spectra of +2, +1, -1 and -2 (iCy3)_2_ Dimer-Labeled ss-dsDNA Constructs

In Table 2, we summarize the parameters obtained for the +2, +1, -1 and -2 (iCy3)_2_ dimer-labeled ss-dsDNA constructs, in which the roll angle *η*_*AB*_ was varied symmetrically. In Table 3, we present a similar summary for anti-symmetric rotation of *η*_*AB*_. The TQ, ED and PD models provide roughly equivalent results for the four labeling positions. The optimized values of the twist angle are very similar among the three models, with ‘right-handed’ conformations (*ϕ*_*AB*_ < 90°) at positive integer positions (∼85° at +2 and +1) and ‘left-handed’ conformations (*ϕ*_*AB*_ > 90°) at negative integer positions (∼96° at -1 and ∼98° at -2). These findings are consistent with previously reported conformations for the same (iCy3)_2_ dimer-labeled ss-dsDNA constructs [14, 15].

**Table 2.** Optimized conformational parameters for (iCy3)_2_ dimer-labeled ss-dsDNA constructs at 25°C based on the TQ, ED and PD models and van der Waals radius *r*_*w*_ = 1.5Å. For these calculations, the roll angle, *η*_*AB*_, was varied symmetrically. Error bars were calculated based on a 1% deviation of the χ^2^ function from its minimum (optimized) value.

| Position | Model (Symm.) | $\phi_{AB}$ (°) | $\theta_{AB}$ (°) | $\eta_{AB}$ (°) | $\delta_{AB}$ (Å) | $\xi_{AB}$ (Å) | $R_{AB}$ (Å) | $\sigma$ (cm<sup>-1</sup>) | $J$ (cm<sup>-1</sup>) |
| --- | --- | --- | --- | --- | --- | --- | --- | --- | --- |
| +2 | TQ | 85±1 | 1±1 | 110±1 | -0.2±0.8 | 2.9±0.2 | 6.2±0.1 | 298±12 | 439 |
|  | ED | 85±1 | 3±15 | 106±40 | 1.6±0.1 | 0.4±1.0 | 7.4±0.1 | 298±12 | 439 |
|  | PD | 84±1 | 10±1 | 124±30 | -0.4±1.0 | -1.5±0.1 | 7.5±0.1 | 298±11 | 439 |
| +1 | TQ | 85±1 | 30±1 | 124±2 | 0.0±0.3 | -3.0±0.1 | 7.3±0.1 | 301±11 | 421 |
|  | ED | 89±1 | 9±3 | 82±60 | 2.0±0.1 | -0.7±0.2 | 7.7±0.1 | 301±11 | 421 |
|  | PD | 89±1 | 13±1 | 111±30 | -1.2±0.1 | -1.7±0.0 | 7.1±0.1 | 302±12 | 422 |
| -1 | TQ | 96±1 | 25±6 | 111±3 | -3.7±0.2 | 0.4±1.0 | 7.0±0.1 | 317±12 | -359 |
|  | ED | 97±1 | 17±10 | 74±230 | -1.3±0.1 | 1.4±1.0 | 8.6±0.1 | 317±12 | -359 |
|  | PD | 98±1 | 13±13 | 128±90 | -4.0±0.3 | 0.7±0.8 | 9.4±0.1 | 317±12 | -359 |
| -2 | TQ | 98±1 | 15±15 | 116±2 | 3.9±0.2 | -0.4±0.6 | 7.5±0.1 | 312±12 | -362 |
|  | ED | 99±1 | 4±3 | 84±60 | -1.1±0.1 | 1.6±0.2 | 8.0±0.1 | 313±12 | -362 |
|  | PD | 95±1 | 32±1.0 | 107±30 | 2.0±0.1 | 0.8±0.1 | 7.1±0.1 | 313±12 | -362 |

**Table 3.** Optimized conformational parameters for (iCy3)_2_ dimer-labeled ss-dsDNA constructs at 25°C based on the TQ, ED and PD models and van der Waals radius *r*_*w*_ = 1.5Å. For these calculations, the roll angle, *η*_*AB*_, was varied anti-symmetrically. Error bars were calculated based on a 1% deviation of the χ^2^ function from its minimum (optimized) value.

| Position | Model (Anti-symm.) | $\phi_{AB}$ (°) | $\theta_{AB}$ (°) | $\eta_{AB}$ (°) | $\delta_{AB}$ (Å) | $\xi_{AB}$ (Å) | $R_{AB}$ (Å) | $\sigma$ (cm<sup>-1</sup>) | $J$ (cm<sup>-1</sup>) |
| --- | --- | --- | --- | --- | --- | --- | --- | --- | --- |
| +2 | TQ | 81±1 | 29±1 | 113±3 | -2.0±0.1 | -0.4±0.4 | 7.4±0.1 | 298±12 | 439 |
|  | ED | 85±1 | 2.0±3 | 120±70 | -1.1±0.5 | 1.8±0.1 | 7.9±0.1 | 298±12 | 439 |
|  | PD | 84±1 | 10±1 | 124±20 | -1.5±0.1 | -0.4±0.2 | 7.5±0.1 | 298±12 | 439 |
| +1 | TQ | 86±1 | 28±1 | 113±3 | -2.9±0.1 | -0.1±0.3 | 8.0±0.1 | 301±11 | 421 |
|  | ED | 89±1 | 14±2 | 129±270 | -2.2±0.3 | 2.8±0.2 | 9.8±0.1 | 301±11 | 421 |
|  | PD | 89±1 | 9±1 | 112±30 | -1.7±0.0 | 0.3±0.2 | 7.1±0.0 | 301±11 | 421 |
| -1 | TQ | 94±1 | 33±5 | 117±2 | 2.1±1.4 | 2.8±0.3 | 7.2±0.1 | 317±12 | -359 |
|  | ED | 98±1 | 16±3 | 71±1 | 3.4±0.2 | -2.0±0.1 | 9.7±0.2 | 317±12 | -359 |
|  | PD | 91±1 | 42±4 | 101±50 | 2.6±0.8 | 1.2±0.1 | 7.2±0.1 | 316±12 | -360 |
| -2 | TQ | 99±1 | 3±20 | 130±2 | 0.8±0.4 | 3.4±0.6 | 8.1±0.1 | 312±12 | -362 |
|  | ED | 96±1 | 27±5 | 106±240 | 0.8±0.4 | -1.7±0.1 | 9.0±0.1 | 312.6±11 | -362 |
|  | PD | 96±1 | 27±7 | 102±60 | 1.8±0.6 | 1.4±0.1 | 7.5±0.1 | 312±12 | -362 |

The agreement between the three models is somewhat less favorable for the tilt, roll and separation parameters. For the +2 position, the TQ model provides a relatively small value for the tilt angle *θ*_*AB*_ = ∼1.2°, while the ED and PD models lead to slightly larger values. As the probe labeling position is varied across the ss-dsDNA junction towards the ssDNA sequences, the TQ model shows an abrupt increase in the tilt angle to ∼30° at the +1 position, which decreases slightly to ∼25° at the -1 position, followed by a decrease to ∼15° at the -2 position. Like the +15 labeled DNA construct, the roll angle adopts a pseudo-face-to-face conformation for all the ss-dsDNA constructs labeled near the fork junction. The value of the roll angle near the ss-dsDNA junction (*η*_*AB*_ = ∼120°) is only slightly more open than that found deep in the DNA duplex region (*η*_*AB*_ = ∼105°). Finally, the inter-chromophore separation is slightly larger at the +2 position (*R*_*AB*_ = ∼6.2Å) than in the duplex region (∼5.9Å). Using parameters obtained with the TQ model, we find that this value for *R*_*AB*_ increases slightly from 6.2Å, 7.3Å, 7.0Å and 7.5Å as the probe labeling position is moved across the ss-dsDNA junction from the +2 to the -2 position.

In Fig. 11, we compare the experimental and simulated absorbance and CD spectra for the +2, +1, -1 and -2 (iCy3)_2_ dimer-labeled ss-dsDNA constructs (panels *A* – *D*, respectively). The *ϕ*_*AB*_, *θ*_*AB*_-dependent χ^2^ surfaces shown correspond to fitting the TQ model to experimental absorbance and CD spectra, where the globally optimized solutions are indicated by red dots. The optimized (iCy3)_2_ dimer conformations are represented schematically as green ball-and-stick models with blue mesh van der Waals surfaces. The agreement between the experimental and simulated absorbance and CD spectra is very good for the +2 (Fig. 11*A*), +1 (Fig. 11*B*) and -1 (Fig. 11*C*) labeling positions. However, the agreement is notably less favorable for the -2 position, especially considering the CD spectrum (Fig. 11*D*). This is perhaps due to the presence of two or more conformational macrostates in solution, with each species contributing a Boltzmann-weighted term to the overall CD signal.

**Figure 11.**
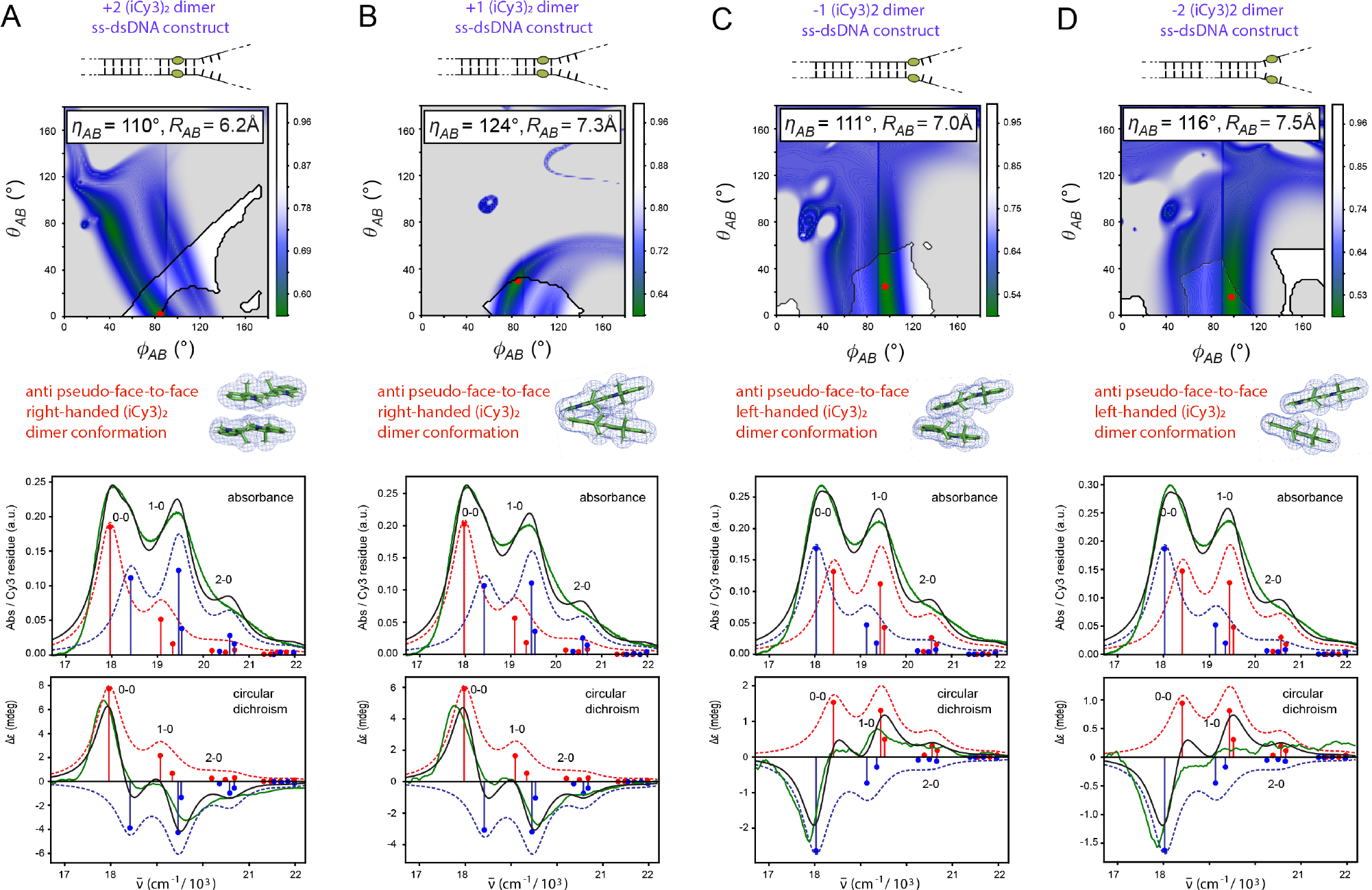
Results of TQ model analysis of experimental absorbance and CD data for the (*A*) +2, (*B*) +1, (*C*) -1 and (*D*) -2 anti (iCy3)_2_ dimer-labeled ss-dsDNA constructs. The top row shows the χ^2^ error function plotted as a function of *ϕ*_*AB*_ and *θ*_*AB*_ for fixed values of *η*_*AB*_, *δ*_*AB*_, *ξ*_*AB*_ and *R*_*AB*_ (see Table 1 for symmetric roll and Table 2 for antisymmetric roll). Semi-transparent gray regions indicate sterically-disallowed conformations using the van der Waals radius *r*_*w*_ = 1.5Å. The red dots indicate the globally minimized values of χ^2^. The second row shows side views of the optimized anti (iCy3)_2_ dimer conformations. Third and fourth rows compare simulated and experimental absorbance and CD spectra, respectively. Additional explanation is provided in the Fig. 10 caption.

In contrast, the (iCy3)_2_ dimer-labeled ss-dsDNA constructs labeled at the +15, +2, +1 and -1 positions are each well-modeled using as a single average dimer conformation, which is consistent with previous findings that these systems exhibit only a moderate level of conformational disorder [15]. We posit that for all the (iCy3)_2_ dimer-labeled ss-dsDNA constructs that we studied, only a small number of dimer conformations are possible due to steric constraints imposed by the surrounding DNA bases and sugar-phosphate backbones. In this picture, for the +15, +2, +1 and -1 dimer-labeled ss-dsDNA constructs, the distribution of conformations is dominated by one of the possible macrostates, while for the -2 labeled DNA construct the distribution likely contains two or more macrostates of near equal stability. More detailed information about the distribution of conformational macrostates under equilibrium conditions can be obtained from single-molecule fluorescence experiments on these same systems [11].

In Fig. 12, we present visualizations of the optimized anti-face-to-face (iCy3)_2_ dimer conformations built into the DNA frameworks of the +2, +1 and -1 DNA constructs. In building these models, we have assumed that the secondary structures of the surrounding DNA bases and backbones are not significantly altered by the presence of the dimer probes. This assumption is borne out by the relatively continuous and non-disrupted conformations of the DNA shown adjacent to these probe positions in Fig. 12. Clearly the conformation of the (iCy3)_2_ dimer probe at a position close to the ss-dsDNA fork junction reflects the local secondary structure of the DNA bases and sugar-phosphate backbones immediately surrounding the dimer probe. To quantify these structural properties, we calculated for each of the dimer-labeled ss-dsDNA constructs the inter-base-pair separation and helical rotation of base pairs above and below the insertion sites of the (iCy3)_2_ dimer probes, which we determined using line segments connecting the C1′ carbons of the glycosidic bonds between complementary bases. At the +2 position, the right-handed (iCy3)_2_ dimer conformation – with inter-base-pair spacing ∼16Å (equal to the spacing of ∼5 stacked bases) and helical rotation ∼19° (equal to the rotation of 0.6 base pairs) – is structurally like that of the +15 dimer-labeled DNA construct (with inter-base-pair spacing ∼16Å and helical rotation ∼56°, see Fig. 10*E*). However, at the +1 position the junction is splayed open and significantly more twisted with inter-base-pair spacing ∼18Å and helical rotation ∼160°. At the -1 position, the opening of the ss-dsDNA fork junction is slightly less pronounced in comparison to the +1 position with helical rotation ∼141°. The position-dependent base-pair spacing and helical rotation introduced by the (iCy3)_2_ dimer probes near the ss-dsDNA junction is likely due to partial unstacking of the Watson-Crick base pairs immediately adjacent to the probes. This decline in base-stacking stability near the ss-dsDNA fork junction is due to the absence of the cooperative stacking interactions present in the adjacent DNA duplex regions, and this effect is likely to be highly sensitive to local base-sequence context.

**Figure 12.**
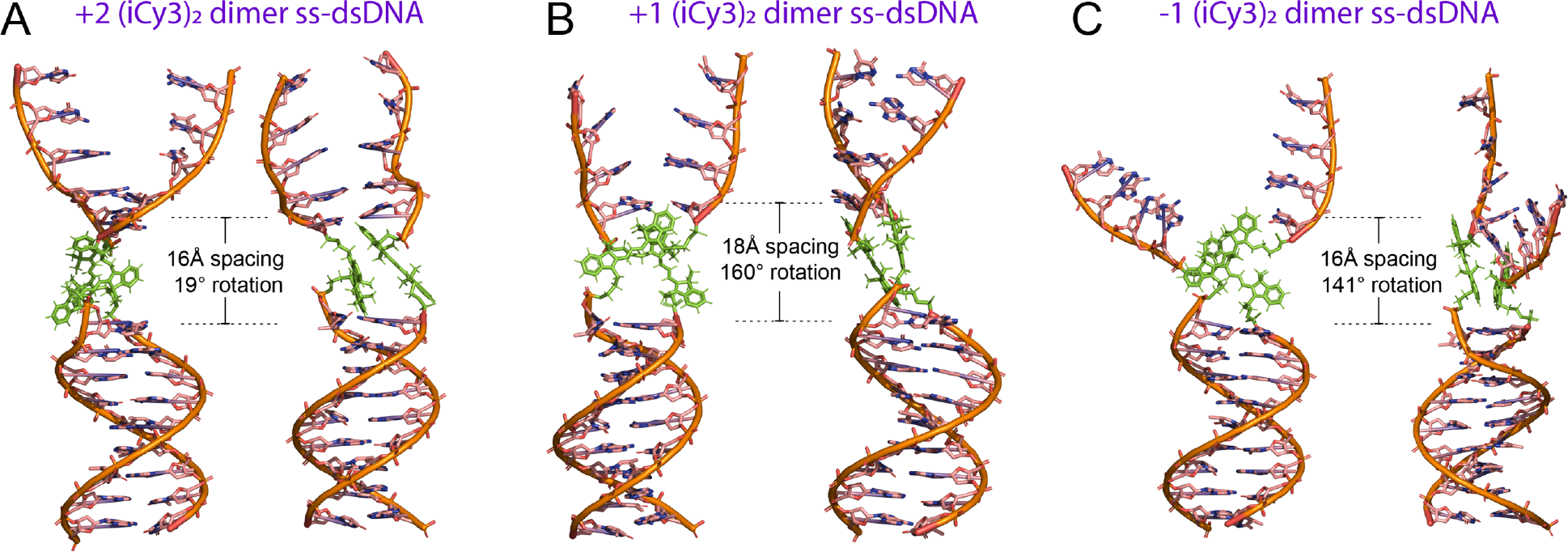
PyMOL visualizations of the optimized anti-face-to-face (iCy3)_2_ dimer conformation built into the sugar-phosphate backbones of the (*A*) +2, (*B*) +1 and (*C*) -1 ss-dsDNA fork constructs. Each structure was geometrically optimized by fixing coordinates of the (iCy3)_2_ dimer atoms and the adjacent bases and backbones using Avogadro (see Sect. II.G). In each panel, the two models are shown rotated 90° relative to one another.

## IV. Discussion

Applying accurate theoretical models for the electrostatic couplings within a molecular dimer is a key step for simulating its optical properties and analyzing its optical spectra. Exciton-coupled (iCy3)_2_ dimer probes, which are rigidly inserted into the sugar-phosphate backbones of ss-dsDNA fork constructs, exhibit absorbance and circular dichroism (CD) spectra that depend on the insertion site position of the dimer probe relative to the ss-dsDNA fork junction. The absorbance and CD spectra of (iCy3)_2_ dimer-labeled ss-dsDNA constructs are generally well-described using a single-effective-mode Holstein-Frenkel Hamiltonian, which includes a conformation-dependent electrostatic coupling parameter, *J*. Values for *J* can be estimated using relatively simple transition density models of the constituent iCy3 monomers. These include the point-dipole (PD), extended-dipole (ED) and transition charge (TQ) electrostatic coupling models, each of which represents a different level of coarse-grained approximation to the monomer transition density. The TQ model represents the highest level of approximation of the transition density, as it more accurately accounts for the full dimensionality of the iCy3 chromophore by assigning individual trESP transition charges to the atomic coordinates.

By implementing a multi-parameter optimization procedure to simulate absorbance and CD spectra, we determined structural coordinates of the (iCy3)_2_ dimer-labeled ss-dsDNA constructs using the PD, ED and TQ models, as summarized in Tables 1 – 3. The conformational parameters that we obtained provide complementary and comparable information about the (iCy3)_2_ dimer-labeled DNA fork constructs. Specifically, the three models produced nearly identical results for the twist angle, *ϕ*_*AB*_. However, the atomistically detailed TQ model can be used to better distinguish tilt, *θ*_*AB*_, roll, *η*_*AB*_, and separation, *R*_*AB*_. To obtain this level of consistency it was necessary to consider steric interactions, since the different levels of approximation otherwise led to diverging results for small inter-chromophore separations.

For the +15 (iCy3)_2_ dimer-labeled ss-dsDNA construct, we obtained an optimized dimer conformation with structural parameters: *ϕ*_*AB*_ = ∼80°, *θ*_*AB*_ = ∼2.4°, *η*_*AB*_ = ∼105° and *R*_*AB*_ = ∼5.9Å. By incorporating the optimized conformation into a DNA structural model (see Fig. 10*E*), we estimate an inter-base-pair spacing of ∼16Å and a helical rotation of 56° that is introduced by the presence of the probes. For the (iCy3)_2_ dimer-labeled ss-dsDNA fork constructs, in which the dimer probes are positioned at and near the ss-dsDNA fork junction (at sites +2, +1, -1 and -2), the roll and separation parameters retain a relatively narrow range of values, independent of position with 110° ≲ *η*_*AB*_ ≲ 124° and 6.2Å ≲ *R*_*AB*_ ≲ 7.5Å. However, the twist and tilt parameters undergo significant changes as the probe position is varied across the ss-dsDNA fork junction towards the single-stranded DNA regions (see Fig. 12). At the +2 position, the conformation is like that at the +15 position with *ϕ*_*AB*_ = ∼85° and *θ*_*AB*_ = ∼1.2°, which gives rise to an inter-base-pair spacing of ∼16Å and a helical rotation of 19°. At the +1 position, the twist angle does not change while the tilt angle increases significantly with *ϕ*_*AB*_ = ∼85° and *θ*_*AB*_ = ∼30°, leading to an inter-base-pair spacing of ∼18Å and a dramatic helical rotation of 160°. At the -1 position, the twist angle increases further to *ϕ*_*AB*_ = ∼96°, and the tilt angle retains its preceding value *θ*_*AB*_ = ∼25°, which gives rise to an inter-base-pair spacing of ∼16Å and a helical rotation of 141°. Our analysis of the −2 (iCy3)_2_ dimer-labeled ss-dsDNA construct did not yield good agreement between simulated and experimental spectra. Because the -2 dimer-labeled ss-dsDNA fork construct could not be well represented using a single average conformation, this system likely exists as a mixture of two or more comparably weighted conformational species in solution.

The results of the current studies provide a basis for the interpretation of experiments that use (iCy3)_2_ dimer-labeled ss-dsDNA constructs to probe protein-DNA binding mechanisms. For example, replication proteins that bind and function at positions near ss-dsDNA fork junctions are likely influenced by DNA ‘breathing’ fluctuations at these sites [1, 11, 20, 30]. Similarly, regulatory proteins that interact with the sugar-phosphate backbones of dsDNA in a non-base-sequence-specific manner likely encounter DNA breathing fluctuations as they undergo a diffusive search process before binding to their specific target sequences [48, 49]. By understanding the local conformations and conformational fluctuations of (iCy3)_2_ dimer-labeled ss-dsDNA constructs, it should be possible to employ these probes in future studies – both at the ensemble and single-molecule level – to gain additional important molecular insights into the detailed mechanisms of biologically significant protein-DNA interactions.

## Data Availability

The data underlying this article are available in the article.

## Acknowledgements

The authors are grateful to their laboratory colleagues for many helpful discussions. K.A.K. wishes to thank Penn State’s Institution of Computational and Data Sciences for providing computing resources used in preliminary work for this project. P.H.v.H. is an American Cancer Society Research Professor of Chemistry.

## References

1. Maurer, A., C. S. Albrecht, P. Herbert, D. Heussman, A. Chang, P. H. von Hippel, and A.H. Marcus, Studies of DNA ‘breathing’ by polarization-sweep single-molecule fluorescence microscopy of exciton-coupled (iCy3)_2_dimer-labeled DNA fork constructs. bioRxiv, 2023. https://www.biorxiv.org/content/10.1101/2023.09.27.559819v2.

2. Acton, T.B., R. Xiao, S. Anderson, J. Aramini, W. A. Buchwald, C. Ciccosanti, K. Conover, J. Everett, K. Hamilton, Y. J. Huang, H. Janjua, G. Kornhaber, J. Lau, D. Y. Lee,G. Liu, M. Maglaqui, L. Ma, L. Mao, D. Patel, P. Rossi, S. Sahdev, R. Shastry, G. V. T. Swapna, Y. Tang, S. Tong, D. Wang, L. Zhao, and G. T. Montelione, Preparation of protein samples for NMR structure, function, and small-molecule screening studies. Meth. Enzymol., 2011. 493: p. 21–60.

3. Dessau, M.A., Y. Modis, Protein crystallization for X-ray crystallography. J. Vis. Exp., 2011. 47: p. 2285.

4. Levitus, M., and S. Ranjit, Cyanine dyes in biophysical research: The photophysics of polymethine fluorescent dyes in biomolecular environments. Quart. Revs. Biophys., 2011. 44(1): p. 123–151.

5. Roy, R., S. Hohng, and T. Ha, A practical guide to single-molecule FRET. Nature Meths., 2008. 5: p. 507–16.

6. Lee, W., P. H. von Hippel, and A. H. Marcus, Internally labeled Cy3 / Cy5 DNA constructs show greatly enhanced photostability in single-molecule FRET experiments. Nucl. Acids Res., 2014. 42: p. 5967–5977.

7. Algar, W.R., N. Hildebrandt, S. S. Vogel, and I. L. Medintz, FRET as a biomolecular research tool — understanding its potential while avoiding pitfalls. Nat. Meth., 2019. 16: p. 815–829.

8. Screenivasan, R., I. A. Shkel, M. Chhabra, A. Drennan, S. Heitkamp, H.-C. Wang, M. A. Sridevi, D. Plaskon, C. McNerney, K. Callies, C. K. Cimperman, and M. T. Record Jr., Fluorescence-detected conformational changes in duplex DNA in open complex formation by Escherichia coli RNA polymerase: Upstream wrapping and downstream bending precede clamp opening and insertion of the downstream duplex. Biochemistry, 2020. 59: p. 1565–1581.

9. Murphy, M.C., I. Rasnik, W. Cheng, T. M. Lohman, and T. Ha, Probing single-stranded DNA conformational flexibility using fluorescence spectroscopy. Biophys. J., 2004. 86: p. 2530–2537.

10. Israels, B., C.S. Albrecht, A. Dang, M. Barney, P.H. von Hippel, and A.H. Marcus, Submillisecond conformational transitions of single-stranded DNA lattices by photon correlation single-molecule FRET. J. Phys. Chem. B, 2021. 125: p. 9426–9440.

11. Phelps, C., W. Lee, D. Jose, P. H. von Hippel, and A. H. Marcus, Single-molecule FRET and linear dichroism studies of DNA ‘breathing’ and helicase binding at replication fork junctions. Proc. Nat. Acad. Sci. USA, 2013. 110: p. 17320–17325.

12. Cunningham, P.D., Y. C. Kim, S. A. Díaz, S. Buckhout-White, D. Mathur, I. L. Medintz, and J. S. Melinger, Optical properties of vibronically coupled Cy3 dimers on DNA scaffolds. J. Phys. Chem. B, 2018. 122: p. 5020–5029.

13. Kringle, L., N. Sawaya, J. R. Widom, C. Adams, M. G. Raymer, A. Aspuru-Guzik, and A. H. Marcus, Temperature-dependent conformations of exciton-coupled Cy3 dimers in double-stranded DNA. J. Chem. Phys., 2018. 148.

14. Heussman, D., J. Kittell, L. Kringle, A. Tamimi, P. H. von Hippel, and A. H. Marcus, Measuring local conformations and conformational disorder of (Cy3)_2_dimers labeled DNA fork junctions using absorbance, circular dichroism and two-dimensional fluorescence spectroscopy. Faraday Disc., 2019. 216: p. 211–235.

15. Heussman, D., J. Kittell, P. H. von Hippel, and A. H. Marcus, Temperature-dependent local conformations and conformational distributions of cyanine dimer labeled singlestranded—double-stranded DNA junctions by 2D fluorescence spectroscopy. J. Chem. Phys., 2022. 156: p. 045101.

16. Cannon, B.L., D. L. Kellis, L. K. Patten, P. H. Davis, J. Lee, E. Graugnard, B. Yurke, and W. B. Knowlton, Coherent exciton delocalization in a two-state DNA-templated dye aggregate system. J. Phys. Chem. A, 2017. 121: p. 6905–6916.

17. Sohail, S.H., J. P. Otto, P. D. Cunningham, Y. C. Kim, R. E. Wood, M. A. Allodi, J. S. Higgins, J. S. Melinger and G. S. Engel, DNA scaffold supports long-lived vibronic coherence in an indocyanine (Cy5) dimer. Chem. Sci., 2020. 11: p. 8546.

18. Hart, S.M., X. Wang, J. Guo, M. Bathe and G. S. Schlau-Cohen, Tuning optical absorption and emission using strongly coupled dimers in programmable DNA scaffolds. J. Phys. Chem. Lett., 2022. 13: p. 1863–1871.

19. Huff, J.S., D. B. Turner, O. A. Mass, L. K. Patten, C. K. Wilson, S. K. Roy, M. S. Barclay, B. Yurke, W. B. Knowlton, P. H. Davis and R. D. Pensack, Excited-state lifetimes of DNA-templated cyanine dimer, trimer, and tetramer aggregates: The role of excitondelocalization, dye separation, and DNA heterogeneity. J. Phys. Chem. B, 2021. 125: p. 10240–10259.

20. Marcus, A.H., D. Heussman, J. Maurer, C. S. Albrecht, P. Herbert, and P. H. von Hippel, Studies of local DNA backbone conformation and conformational disorder using sitespecific exciton-coupled dimer probe spectroscopy. Ann. Rev. Phys. Chem., 2023. 74: p. 245–65.

21. Sorour, M., A. H. Marcus, and S. Matsika, Modeling the electronic absorption spectra of the indocarbocyanine Cy3. Molecules, 2022. 27: p. 4062.

22. Halpin, A., P. J. M. Johnson, R. Tempelaar, R. S. Murphy, J. Knoester, T. L. C. Jansen, and R. J. D. Miller, Two-dimensional spectroscopy of a molecular dimer unviels the effects of vibronic coupling on exciton coherences. Nature Chem., 2014. 6: p. 196–201.

23. Duan, H.-G., P. Nalbach, V. I. Prokhorenko, S. Mukamel, and M. Thorwart, On the origin of oscillations in two-dimensional spectra of excitonically-coupled molecular systems. New Journal of Physics, 2015. 17: p. 072002.

24. Hestand, N.J., and F. C. Spano, Expanded theory of H- and J-molecular aggregates: The effects of vibronic coupling and intermolecular charge transfer. Chem. Revs., 2018. 118: p. 7069–7163.

25. Kistler, K.A., C. M. Pochas, H. Yamagata, S. Matsika, and F. C. Spano, Absorption, circular dichroism, and photoluminescence in perylene diimide bichromophores: polarization-dependent H- and J-aggregate behavior. J. Phys. Chem. B, 2011. 116: p. 77–86.

26. Kistler, K.A., F. C. Spano, and S. Matsika, A benchmark of excitonic couplings derived from atomic transition charges. J. Phys. Chem. B, 2013. 117: p. 2032–2044.

27. Sorour, M.I., K. A. Kistler, A. H. Marcus, and S. Matsika, Accurate modeling of excitonc coupling in cyanine dye Cy3. J. Phys. Chem. A, 2021. 125: p. 7852–7866.

28. Ahmad, M.F., N. A. M. Isa, W. H. Lim, K. M. Ang, Differential evolution: A recent review based on state-of-the-art works. Alexandria Eng. J., 2022. 61: p. 3831–3872.

29. Storn, R., and K. Price, Differential evolution — A simple and efficient heuristic for global optimization over continuous spaces. J. Global Optim., 1997. 11: p. 341–359.

30. Lee, W., D. Jose, C. Phelps, A. H. Marcus, and P. H. von Hippel, A single-molecule view of the assembly pathway, subunit stoichiometry and unwinding activity of the bacteriophage T4 primosome (helicase-primase) complex. Biochemistry, 2013. 52: p. 3157–3170.

31. Jia, K., Y. Wan, A. Xia, S. Li, F. Gong, and G. Yang, Characterization of photoinduced isomerization and intersystem crossing of the cyanine dye Cy3. J. Phys. Chem. A, 2007. 111: p. 1593–1597.

32. Aydin, M., Ö. Dede, and D. L. Akins, Density functional theory and Raman spectroscopy applied to structure and vibrational mode analysis of 1,1’,3,3’-tetraethyl-5,5’,6,6’tetrachloro-benzimidazolocarbocyanine iodide and its aggregate. J. Chem. Phys., 2011. 134: p. 064325–1-12.

33. Bässler, H., B. Schweitzer, Site-selective fluorescence spectroscopy of conjugated polymers and oligomers. Acc. Chem. Res., 1999. 32: p. 173–182.

34. Valcunas, L., D. Abramavicius, and T. Mančal, Molecular Excitation Dynamics and Relaxation: Quantum Theory and Spectroscopy. 2013: Wiley-VCH.

35. Czikklely, V., H. D. Forsterling, and H. Kuhn, Extended dipole model for aggregates of dye molecules. Chem. Phys. Lett., 1970. 6(3): p. 207–210.

36. Didraga, C., A. Pugžlys, P. R. Hania, H. von Berlepsch, K. Duppen, and J. Knoester, Structure, spectroscopy, and microscopic model of tubular carbocyanine dye aggregates. J. Phys. Chem. B, 2004. 108: p. 14976–14985.

37. Howard, I.A., F. Zutterman, G. Deroover, D. Lamoen, and C. Van Alsenoy, Approaches to calculation of exciton interaction energies for a molecular dimer. J. Phys. Chem. B, 2004. 108: p. 19155–19162.

38. Virtanen, P., R. Gommers, T. E. Oliphant, M. Haberland, T. Reddy, D. Cournapeau, E. Burovski, P. Peterson, W. Weckesser, J. Bright, S. J. van der Walt, M. Brett, J. Wilson, K. J. Millman, N. Mayorov, A. R. J. Nelson, E. Jones, R. Kern, E. Larson, C. J. Carey, I. Polat, Y. Feng, E. W. Moore, J. VanderPlas, D. Laxalde, J. Perktold, R. Cimrman, I. Henriksen, E.A. Quintero, C.R. Harris, A. M. Archibald, A. H. Ribeiro, F. Pedregosa, P. van Mulbregt, SciPy 1.0: Fundamental algorithms for scientific computing in Python. Nat. Meth., 2020. 17: p. 261–272.

39. Rowland, R.S., and R. Taylor, Intermolecular nonbonded contact distances in organic crystal structures: Comparison with distances expected from van der Waals radii. J. Phys. Chem., 1996. 100: p. 7384–7391.

40. Lu, T., F. Chen, Multiwfn: a multifunctional wavefunction analyzer. J. Comput. Chem., 2012. 33: p. 580–592.

41. Breneman, C.M., K. B. Wiberg, Determining atom-centered monopoles from molecular electrostatic potentials. The need for high sampling density in formamide conformational analysis. J. Comput. Chem., 1990. 11: p. 361–373.

42. The PyMOL Molecular Graphics System. 2020, Schrödinger, LLC.

43. Hanwell, M.D., D. E. Curtis, D. C. Lonie, T. Vandermeersch, E. Zurek and G. R. Hutchison, Avogadro: an advanced semantic chemical editor, visualization, and analysis platform. J. Cheminformatics, 2012. 4(17).

44. Hestand, N.J., and F. C. Spano, Interference between Coulombic and CT-mediated couplings in molecular aggregates: Hto J-aggregate transformation in perylene-based πstacks. J. Chem. Phys., 2015. 143: p. 244707.

45. Sinden, R.R., DNA Structure and Function. 1994, San Diego: Academic Press.

46. Datta, K., N. P. Johnson, G. Villani, A. H. Marcus, and P. H. von Hippel, Characterization of the 6-methyl isoxanthopterin (6-MI) base analog dimer, a spectroscopic probe for monitoring guanine base conformations at specific sites in nucleic acids. Nucl. Acids Res., 2012. 40(3): p. 1191–202.

47. Jose, D., K. Datta, N. P. Johnson, and P. H. von Hippel, Spectroscopic studies of position-specific DNA “breathing” fluctuations at replication forks and primer-template junctions. Proc Natl Acad Sci U S A, 2009. 106(11): p. 4231–4236.

48. von Hippel, P.H., From “simple” DNA-protein interactions to the macromolecular machines of gene expression. Annu. Rev. Biophys. Biomol. Struct., 2007. 36: p. 79–105.

49. von Hippel, P.H., N. P. Johnson, and A. H. Marcus, 50 years of DNA ‘breathing’: Reflections on old and new approaches. Biopolymers, 2013. 99: p. 923–954.

